# The solute carrier superfamily interactome

**DOI:** 10.1101/2024.09.30.615192

**Authors:** Fabian Frommelt, Rene Ladurner, Ulrich Goldmann, Gernot Wolf, Alvaro Ingles-Prieto, Eva Lineiro-Retes, Zuzana Gelová, Ann-Katrin Hopp, Eirini Christodoulaki, Shao Thing Teoh, Philipp Leippe, Manuele Rebsamen, Sabrina Lindinger, Iciar Serrano, Svenja Onstein, Christoph Klimek, Barbara Barbosa, Anastasiia Pantielieieva, Vojtech Dvorak, J. Thomas Hannich, Julian Schoenbett, Gilles Sansig, Tamara A.M. Mocking, Jasper F. Ooms, Adriaan P. IJzerman, Laura H. Heitman, Peter Sykacek, Juergen Reinhardt, André C Müller, Tabea Wiedmer, Giulio Superti-Furga

## Abstract

Solute carrier (SLC) transporters form a protein superfamily that enables transmembrane transport of diverse substrates including nutrients, ions and drugs. There are about 450 different SLCs, residing in a variety of subcellular membranes. Loss-of-function of an unusually high proportion of SLC transporters is genetically associated with a plethora of human diseases, making SLCs a rapidly emerging but challenging drug target class. Knowledge of their protein environment may elucidate the molecular basis for their functional integration with metabolic and cellular pathways and help conceive pharmacological interventions based on modulating proteostatic regulation. We aimed at obtaining a global survey of the SLC protein interaction landscape and mapped the protein-protein interactions of 396 SLCs by interaction proteomics. We employed a functional assessment based on RNA interference of interactors in combination with measurement of protein stability and localization. As an example, we detail the role of a SLC16A6 phospho-degron, and the contributions of PDZ-domain proteins LIN7C and MPP1 to the trafficking of SLC43A2. Overall, our work offers a resource for SLC-protein interactions for the scientific community.

## Introduction

Biological membranes demark organismic, cellular and organellar boundaries and are essential for life. The transport of all sorts of organic and inorganic molecules across membranes by membrane-spanning proteins is regulated at many levels, from transcriptional to post-translational, directed by availability of solutes as well as biosynthetic, metabolic and informational requirements.

Within human cells, it is estimated that between 1,500 and 2,000 genes contribute to transport across membranes (Ye *et al*, 2014). Transporters take a key role to enable homeostasis, growth and proliferation. Solute carrier (SLC) proteins can transport substrates using electrochemical gradients and are the largest transporter class with more than 450 members (Meixner *et al*, 2020; Ferrada & Superti-Furga, 2022; He *et al*, 2009; Haas *et al*, 2014). The SLC superfamily covers a wide range of transported substrates and is differentially expressed across cell types and tissues, and across healthy and disease states (Hediger, 2004; Pizzagalli *et al*, 2021; Lin *et al*, 2015). In fact, 244 of 456 SLCs have a link to human diseases (Kelleher *et al*, 2023) (GlobalData; Goldmann et al.). The targets of some of the most important so-called blockbuster drugs, such as serotonin-uptake inhibitors and gliflozins (glucose re-uptake inhibitors) are solute carriers (Wang *et al*, 2020; César-Razquin *et al*, 2018). Despite the estimated potential of 75% of all SLCs to carry small organic molecules, most chemical efforts have focused on just a few SLC families, leaving most families and most SLCs untargeted (Dvorak & Superti-Furga, 2023; Carter *et al*, 2019; Fauman *et al*, 2011; Galetin *et al*, 2024; Gyimesi & Hediger, 2022; Wang *et al*, 2020). SLCs are increasingly considered attractive novel therapeutic targets for modifying disease by modulating metabolism. Strategies for the development of inhibitors have often started from the natural substrate, like in the case of PF-06649298 and SLC13A5, a sodium-coupled citrate transporter primarily expressed in the liver, brain, and other tissues (Huard *et al*, 2015). In the search for alternative routes, it has also been shown that it is possible to target SLC transporters using Proteolysis Targeting Chimeras (PROTACs), causing efficient degradation of the target (Bensimon *et al*, 2020; Zhang *et al*, 2024). However, the majority of SLC-associated diseases represent loss-of-function situations where mutations impair protein levels, such as SLC6A8 and creatine transport deficiency (Ferrada *et al*, 2024; Valayannopoulos *et al*, 2013) or SLC39A8, the metal transporter associated with congenital disorder of glycosylation type II and Leigh syndrome (Park *et al*, 2015). The success in partially restoring the function of hypomorphic alleles of the cystic fibrosis transmembrane conductance regulator (CTFR) ion channel gene by chemical modulators (correctors and potentiators) highlights the possibility that even partial restoration of protein folding and localization can be of clinical benefit (Liu *et al*, 2024), suggesting an avenue for the hundreds of SLC associated loss-of-function pathologies.

Due to the substantial lack of cellular, molecular and functional data for a large portion of the SLC superfamily (César-Razquin *et al*, 2015), the RESOLUTE consortium, a public-private partnership, used a systematic multi-omics approach to gather molecular and functional data on SLCs (**Fig. 1A**), thus complementing the interaction proteomics approach with parallel investigations in metabolomics, transcriptomics (Wiedmer and Teoh et al, accompanying manuscript), genetic interactions (Wolf and Leippe, accompanying manuscript), and multi-omics data analysis (Goldmann et al, accompanying manuscript). Characterization of the multiprotein complexes to which SLC transporters participate represents an important functional dimension of SLCs because of their potential to reveal connections to the biological and biochemical machinery of the cell. The characterization of the protein interactome of SLCs may inform novel therapeutic approaches (Yan *et al*, 2020; Gomez *et al*, 2023; Boeszoermenyi *et al*, 2023). Further, an interaction proteomics approach can reveal chaperones involved in protein folding, which on top of representing therapeutic intervention routes can improve heterologous expression of SLCs and subsequent structural characterization (Torres *et al*, 2003).

**Figure 1.**
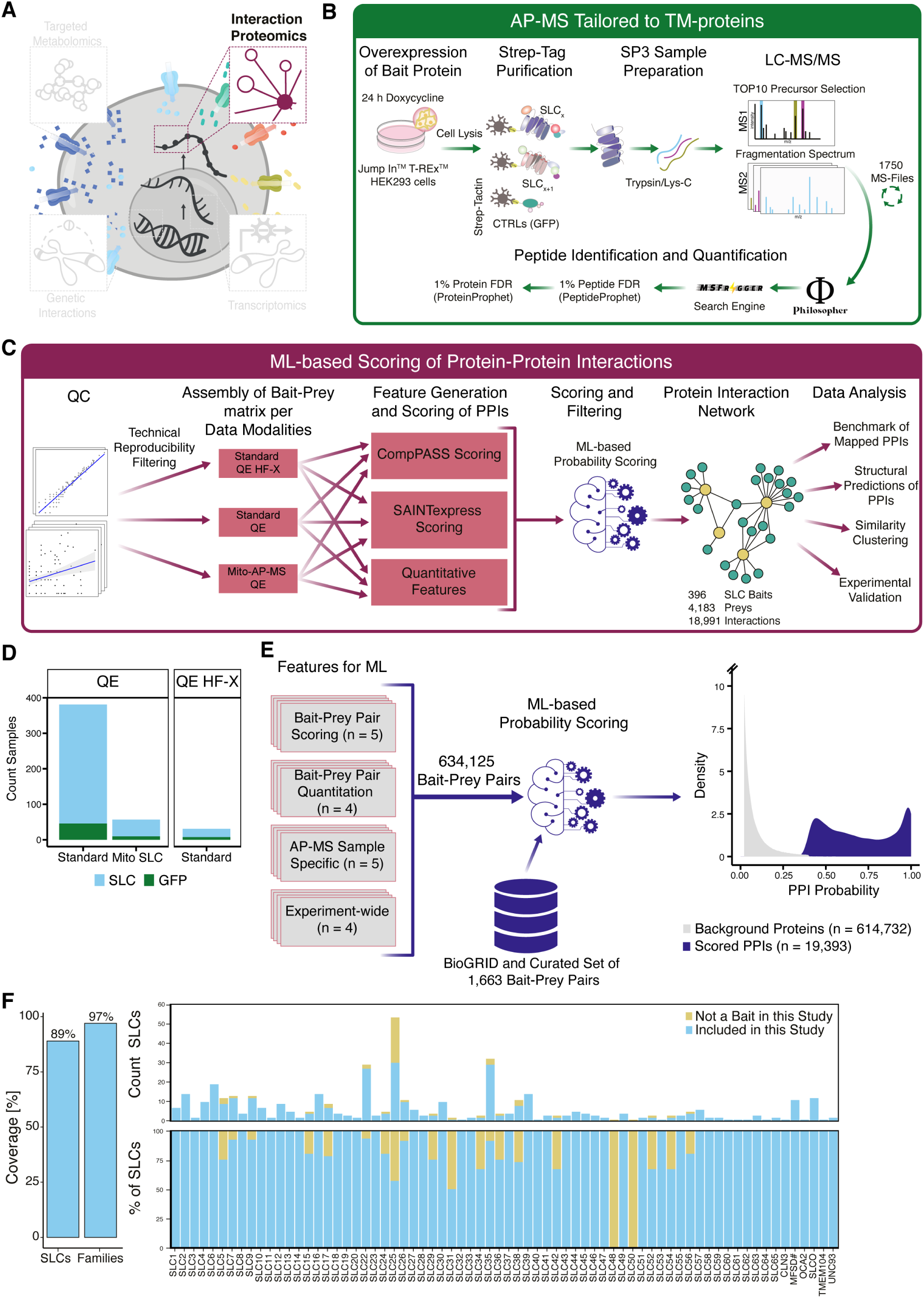
A systematic AP-MS approach to define the human solute carrier superfamily interactome. **(A)** Overview of the - omics layers to characterize SLCs on a molecular level generated within the RESOLUTE project. **(B)** Experimental workflow for native purification of SLC-containing protein complexes from HEK 293 Jump In T-Rex cell lines**. (C)** Modular data analysis pipeline for processing of MS-data and scoring of PPIs. **(D)** SLC baits grouped by protocol and MS-platform. **(E)** A set of 18 features (**Table EV2**) and a curated list of labelled PPIs served as input for the scoring. Distribution of scored interactions (dark blue) versus background proteins (grey). For visualization of both distributions, the y-axis was cut at a density of 10. **(F)** AP-MS coverage across the SLC superfamily. SLCs used as bait in the SLC-interactome are marked in blue, SLCs not included in this study are coloured in yellow.

The protein interactome plays a crucial role for all cell activities, but the interactions of membrane proteins are difficult to study by standard biochemical protein interaction techniques (Schey *et al*, 2013; Cao *et al*, 2021). Human proteome wide interactome studies using mass spectrometry have therefore only reported on a subset of transmembrane domain containing proteins (TM-proteins) and of SLCs in particular, often with protocols developed to mainly suit the study of soluble proteins (Huttlin *et al*, 2021, 2015, 2017). Isolated interactome analyses of individual TM-proteins, on the other hand, can miss the necessary context to understand if interactions are specific or are common cellular interaction modalities for multiple-pass TM-proteins localised on all kinds of subcellular membranes. This is a particular challenge for highly sensitive proteomics, for which several frameworks were developed to assign pairwise protein interaction confidence across large-scale affinity-purification coupled to mass spectrometry (AP-MS) data sets (Teo *et al*, 2014; Sowa *et al*, 2009). AP-MS, as a state-of-the-art method to assign protein interactions offers high sensitivity and reproducibility (Varjosalo *et al*, 2013), and thus is especially suited for large-scale studies (Gavin *et al*, 2002; Huttlin *et al*, 2021; Uliana *et al*, 2023; Buljan *et al*, 2020; Salokas *et al*, 2022).

Studies which focused on single SLCs established the general applicability of MS-based interaction proteomics and linked two transporters to important cellular metabolic machineries. Among these are the interaction of the amino acid transporter SLC38A9 with the Ragulator/LAMTOR complex (Rebsamen *et al*, 2015; Wang *et al*, 2015) and the interaction of SLC15A4 with TASL (Heinz *et al*, 2020). Interaction proteomics studies have also revealed SLC heterodimers, including SLC16A family members with BSG/EMB (Howard *et al*, 2010; Rusu *et al*, 2017), SLC7A family members with SLC3A1/SLC3A2 (Yan *et al*, 2020; Parker *et al*, 2021) and within members of the ZIP-family of Zinc transporters (SLC39A-family) (Taylor *et al*, 2016). Other examples showed that protein interactions determine the accurate folding and ER export of SLCs through chaperone interactions (Wiktor *et al*, 2021; Ohtsubo *et al*, 2011) or affect the correct trafficking of SLCs (Yang *et al*, 2018; El-Kasaby *et al*, 2010). In addition, several SLCs are regulated through proteostatic pathways (Colaco *et al*, 2023; Xu *et al*, 2016) or endocytosis (Puris *et al*, 2022; Rotin & Staub, 2012).

Here, we used a systematic AP-MS approach to determine the protein interaction network of human solute carriers. For this we isolated and analysed over 400 SLC proteins using improved purification strategies from human embryonic kidney cells. We then compared all individual SLC-interactomes and in combination with machine learning (ML), assembled 18,991 interactions for 396 individual SLCs spanning 68 different SLC families.

We integrated gene ontology and metabolic information to understand the nature of these novel interactions and identified individual interacting proteins and protein complexes linked to organelle function, protein trafficking, protein degradation, posttranslational modification and signalling. By genetic loss-of-function analysis, we found dozens of interactors that led to a change in protein stability of the interacting transporter, charting the proteostatic regulatory network of SLC transporters.

## Results

### Obtaining a robust protein interactome of SLC transporters

For the systematic characterization of SLC-protein interactions we aimed to establish a robust, scalable experimental affinity enrichment protocol combined with a reproducible MS-based proteomics method suitable to map the entire human interactome of SLCs in a specific cellular setting.

All the 405 SLCs profiled within this study were subjected to an experimental workflow specifically adapted for TM-proteins (**Fig. 1B**), and built on previously described AP-MS protocols (Varjosalo *et al*, 2013; Gavin *et al*, 2002; Rebsamen *et al*, 2015; Uliana *et al*, 2023). Full elution of TM-proteins was achieved by employing an SDS-based elution. For buffer exchange to remove remaining SDS, we adapted an SP3-based sample clean-up protocol (Müller *et al*, 2020), which enabled scaling up sample preparation. For SLCs localized to mitochondria, we performed a crude mitochondrial enrichment prior to lysis of the mitochondrial fraction by digitonin affinity purification (see methods for details). The resulting AP-samples were acquired by data dependent acquisition (DDA) in biological duplicates with technical injections.

To perform AP-MS for the human SLC superfamily, the codon-optimized consensus sequence of each SLC bait protein was either N- or C-terminally fused to a double-strep-tag and HA-epitope (**Table EV1**) and stably expressed under a doxycycline inducible promoter in HEK 293 Jump In T-REx cells. HEK 293 were also used as model cell line for many different large-scale interaction proteomics studies focusing on individual protein families such as RTKs (Salokas *et al*, 2022) or kinases (Varjosalo *et al*, 2013; Buljan *et al*, 2020), and for the human proteome-wide series BioPlex studies (Huttlin *et al*, 2015, 2017, 2021), thus allowing the comparison of the SLC-interactome data to these data sets. A project of this magnitude required a computational pipeline which was able to handle large amounts of MS- injections and integrate a fine-tuned scoring framework to account for differences in expression and subcellular localization of the various SLCs. We set up a modular processing pipeline employing Philosopher (da Veiga Leprevost *et al*, 2020), which we integrated with an in-house developed scoring approach (**Fig.1C**). Our scoring approach considered also the three data modalities, which were obtained by using two different affinity enrichment protocols and two MS-platforms (**Fig. 1D**). For the ML-scoring, a combination of 18 features was derived for all 634,125 interaction pairs (**Fig. 1E, Table EV2**), serving as input for an RBF classifier (methods for details).

In total, we acquired data for 405 SLCs, with 396 SLCs remaining in the interaction proteomics data sets after scoring and filtering of background proteins. The SLC-interactome covered 97% of the SLC families and 89% of the 447 SLCs which were investigated by the RESOLUTE consortium (**Fig. 1F, Dataset EV1**) (Superti-Furga *et al*, 2020).

### General characteristics of the SLC-interactome

We first assessed bait abundance and its influence on total signal quantified within a sample and on the number of interactors (**Supplementary Results and Appendix Fig. S1A-C).** SLC bait abundance did not correlate with the number of interactors or the total signal, indicating no major bias due to expression levels of the tagged SLC. Further, we compared the log2FC of each SLC bait against GFP controls as well as all other AP-MS samples (**Appendix Fig. S1D**). Several SLCs showed lower enrichment against the membrane background of other SLC AP-MS samples than to GFP controls. We concluded that AP-MS data from other SLCs represented a better strategy to correct for co-purified SLCs.

The interactors had a signal that was at least two-fold higher than before filtering (**Appendix Fig. S2A**). Across our data set we quantified on average 1,566 proteins per SLC, indicative of a highly complex membrane background (**Appendix Fig. S2B**), albeit with strong variation (a few hundred to 3,000 proteins). This may be due to SLCs being expressed in different compartments and the fact that the data were obtained using different MS-platforms. We assigned 48 interaction partners, which roughly corresponds to the number of interaction partners when investigating an individual SLC (Rebsamen *et al*, 2015) and to other large scale human interaction proteomics studies (Uliana *et al*, 2023; Sowa *et al*, 2009; Ciuffa *et al*, 2022). A few SLCs retrieved a higher number of interaction partners, maybe reflecting their multiple subcellular localizations (**Appendix Fig. S2C**). To test for a bias towards frequently or uniquely quantified proteins, we plotted the frequency of observation against the quantitative signal of each interactor (**Appendix Fig. S2D**). We scored equal numbers of interactors across the SLC-interactome, suggesting no penalty for frequently quantified proteins.

To determine the network properties of the fully assembled SLC-interactome, we transformed the obtained network into an undirected network graph and calculated the degree of connectivity: the number of adjacent edges per bait or interaction partner. In addition, we calculated the Kleinberg’s hub centrality score, which is a measure of node influence on connecting other nodes (Kleinberg, 1999).

Within the network, around 25% of the nodes were uniquely connected, whereas most nodes were densely connected with a median of 33 edges (**Appendix Fig. S2E**). The top 20 most connected interaction partners of the SLC-interactome included several interactions that were previously linked to TM-proteins and/or SLC function (**Appendix Fig. S2F**). Among these were the folding chaperone calmegin (CLGN) (Huttlin *et al*, 2021), ANKRD13A, an ankyrin repeat domain-containing protein involved in the internalization of receptors (Mattioni *et al*, 2020) and the ER-localized chaperones ERLIN1/2 (Wiktor *et al*, 2021). Investigating the most influential hubs across the binary network revealed 54 interaction partners and 46 SLCs among the 100 most important hubs (**Appendix Fig. S2G**). Among the most connected interactors were proteins involved in the glycosylation machinery (e.g.: DDOST, RPN1/2, STT3A, GALNT2/7) and protein trafficking (e.g.: GOLGA5, SEC62, CNIH4, **Appendix. Fig. S2G**).

### Benchmarking of the SLC-interactome

To determine the relevance of the data, we benchmarked the interaction data on a PPI level against reported PPIs in databases as well as on a multi-subunit protein complex level. Compared to other TM-proteins or proteins without an annotated TM-domain, few SLCs are characterized by AP-MS and fewer PPIs are reported, making benchmarking challenging (**Fig. 2A)**.

**Figure 2.**
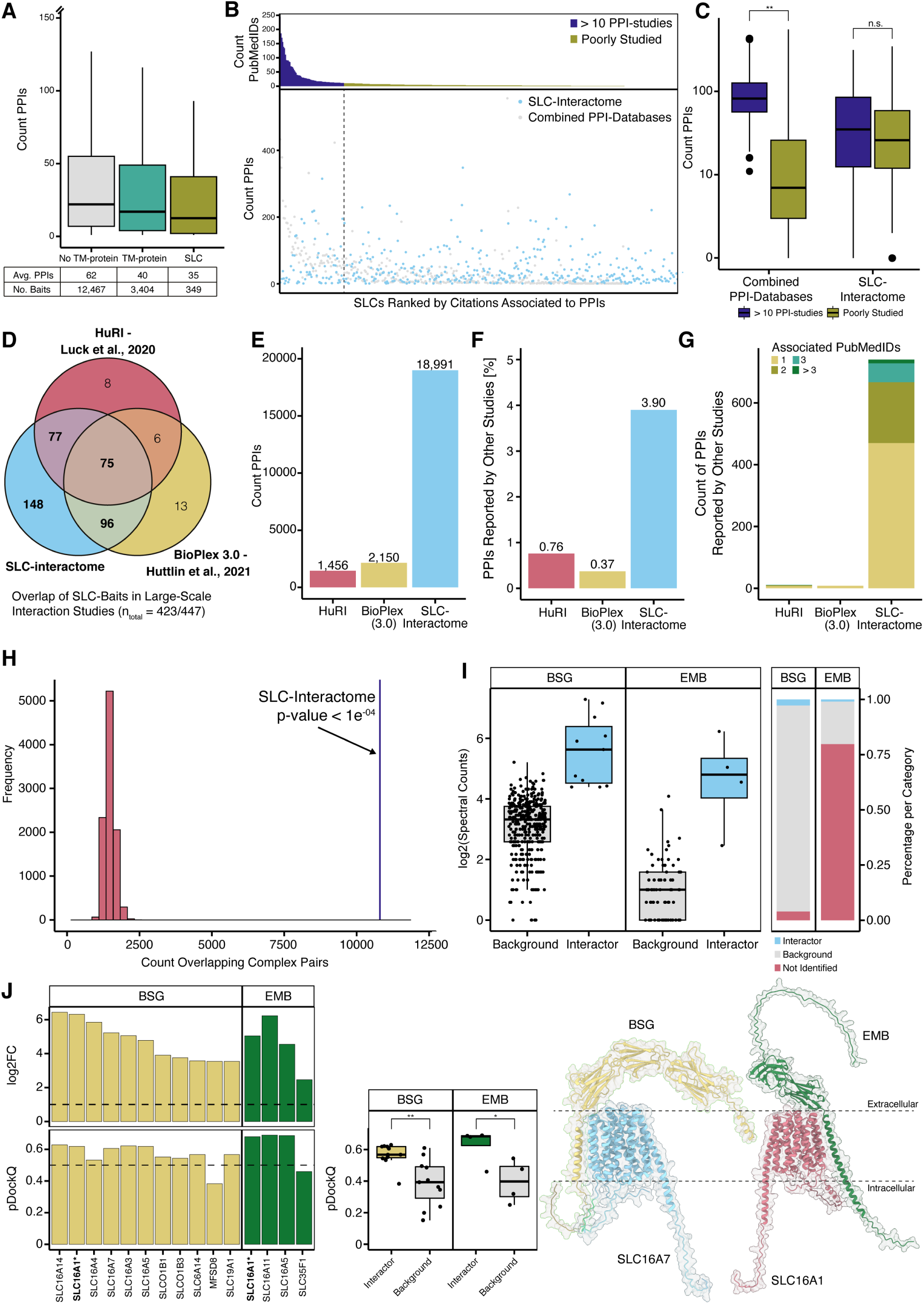
Assessment of SLC-protein interaction network and data quality. **(A)** Reported protein interactions for SLCs (blue) versus TM-domain containing proteins (green) and proteins without annotated TM-domain(grey). **(B)** PPIs per SLC reported in the PPI-library. The upper part shows the associated studies per PPI, and the lower part the number of reported PPIs (grey dots) and PPIs identified within the SLC-interactome (blue dots). SLCs were grouped by the associated publication into a group of SLC which were studied by interaction proteomics (dark blue, > 10 referenced studies) and a group of poorly characterized SLCs (yellow, < 10 associated studies or none). **(C)** Comparison of PPIs reported for the poorly studied (yellow) against more often studied SLCs (dark blue). For the SLC-interactome no bias was observed (in literature poorly characterized, n=332, average PPIs=45.4, > 10 associated studies, n=64, average PPIS=61.1, Wilcox rank sum test p-value=0.3318) in comparison to the literature reported database for which a statistically significant bias between SLCs was found (poorly characterized, n=375, average PPIs=21.8, > 10 associated studies, n=72, average PPIs=111, Wilcox rank sum test p-value< 2.2e-16). **(D)** Overlap of SLCs used as baits in the SLC-interactome study (blue), in the BioPlex (yellow) and HuRI (red). **(E)** PPIs reported for SLCs in the BioPlex (yellow), HuRI and the SLC-interactome (blue). **(F)** Fraction of protein interactions reported by BioPlex (yellow), HuRI (red) and the SLC-interactome (blue) that were reported by additional studies. **(G)** PPIs reported in literature and the two large-scale reference studies and the SLC-interactome. The colour indicates associated studies for reported PPIs. **(H)** CORUM derived interaction pairs are enriched within the SLC-interactome in comparison to 10,000 permutated networks with conserved topology and composition. **(I)** Distribution of BSG and EMB, two chaperones of SLC16A-family members, across the SLC-interactome. The left panel shows the log2 transformed SPCs separated by scored interactions (blue) and background (grey) within the SLC-interactome. Bars on the right side indicate how often the chaperones were scored or found as background across the SLC-interactome. **(J)** Upper part shows for SLC-BSG (yellow) and EMB (green) interactions the log2FC against GFP (dotted black line log2FC > 1), and the lower part shows the pDockQ (dotted black line pDockQ > 0.5, high-confidence structures). Complexes for which the structure was experimentally solved are marked with an asterisk and are in bold (*). Predicted SLC-chaperone structures were compared against a set of predicted structures of SLC-chaperones for which the chaperones were classified as background (unpaired student t-test for BSG p-value=0.001589 and EMB with a p-value=0.02762; independent control sets and tests). On the right-side predicted structures for SLC16A7-BSG and SLC16A1-EMB complexes are shown. For all boxplots in the figure panels: Box represents the interquartile range and its whiskers 1.5 X IQR. Black line represents the median and the black dots represents single measurements.

We generated a combined reference PPI library retrieving interactions from multiple sources including BioGRID (Oughtred *et al*, 2021), IID (Kotlyar *et al*, 2022), IntAct (Orchard *et al*, 2014) and STRING (Szklarczyk *et al*, 2023). We subsequently filtered the PPI-library to contain interactions obtained by AP-MS and equivalent interaction mapping techniques (see methods). The resulting reference PPI- library covered 404 (90%) of the 447 SLCs and a total of 16,072 PPIs (**Fig. EV1A**). The reference PPI- library showed considerable variation with 197 SLCs (44%) and only 1,214 PPIs (7.55%) overlapping across databases and nearly half of the PPIs (7,899) were only reported by a single database, thus highlighting first the need to combine PPI-libraries and secondly to add high-quality SLC interactome data as a new reference point (**Fig. EV1B**).

We compared the SLC-interactome and reference PPI-library using the similarity matrix between the structural models of human SLCs (Ferrada & Superti-Furga, 2022), and represented the relationships by an unrooted structural similarity tree, similar to how the human kinases were shown in the kinome tree (Manning *et al*, 2002; Karaman *et al*, 2008). Unlike the kinome tree, there is no common ancestor, and phylogenetic relationships are valid only within one fold (details in Goldmann et al., accompanying manuscript). With each node representing an individual SLC, the edges were coloured by the major structural folds of the SLCs, and all SLCs included in the study were coloured in blue (**Fig. EV2**). 24 clade classifications with an additional clade classified as unknown were annotated. Clustering and fold classification were as described in Ferrada and Superti-Furga (Ferrada & Superti-Furga, 2022). The SLCome was decorated with the PPIs per clade, scored within the SLC- interactome and reported in the reference PPI-library (see methods). For 23 clades we found 18,232 novel PPIs, increasing the number of PPIs up to 6-fold per clade and indicating that the PPI coverage is mostly independent of structural and evolutionary features.

We found less interactions compared to the reference PPI-library for six structural clades, among them the MitC clade (3,913 in literature vs. 454 within the SLC-interactome) and the SLC56 clade (674 vs. 14). These two clades consist of mitochondrial transporters. This might be because highly abundant mitochondrial carriers are often wrongly assigned as interactors, as suggested by their high CRAPome presence (Mellacheruvu *et al*, 2013). Eleven SLCs from the reference PPI-library had a frequency above 20%, indicating non-specific engagement to the beads-matrix, and of those, eight were mitochondrial SLCs. To investigate if SLCs found with a high CRAPome presence are overrepresented as interaction partners, we calculated the ratio of these SLCs in the bait or prey role for each PPI reported in BioGRID (**Fig. EV1C**). We found that SLCs with higher presence in the CRAPome database were significantly more often reported as interactors (90.24% vs. 60.45%), indicating that many of the interactions were likely non-specific (**Fig. EV1D**).

We next investigated the potential bias of currently reported PPIs towards heavily studied superfamily members, as shown for example for kinases (Buljan *et al*, 2020). As expected, the number of reported interactions and associated studies strongly varied between SLCs (**Fig. 2B**). We grouped the SLCs according to the number of references: more than 10 references (72 SLCs), less than 10 references (315 SLCs) or no associated references (60 SLCs) (**Fig. 2B**). Comparing the number of PPIs between highly studied or poorly characterized SLCs revealed a significant difference for the reference PPI library (Welch two sample t-test p-value 9.791e^-12^), but not the SLC-interactome (Welch two sample t-test p-value 0.0896), thus supporting that our systematic approach provides novelty across the SLC superfamily independent of prior knowledge (**Fig. 2C**).

Comparing the SLC-interactome to other large-scale studies, we found that 148 baits had not been covered by other large-scale AP-MS (Huttlin *et al*, 2021) or yeast-two-hybrid (Luck *et al*, 2020) studies (**Fig. 2D**). This study almost doubles the number of SLC interactomes of previous large-scale studies and contains approximately ten times more PPIs for the SLC superfamily (**Fig. 2E**). When comparing all three data sets against the reference PPI-library, the SLC-interactome had, with 3,9%, the highest recovery rate (**Fig. 2F**) and identified the largest number of PPIs with two or more associated studies (**Fig. 2G**). Within our data set we found 740 PPIs that were previously identified, compared to 11 and 8 for HuRI and BioPlex, respectively. A total of 96.1% interactions within the SLC-interactome are novel, which is slightly higher than comparable interaction proteomics studies that identified 80 to 85% novel interactions (Taipale *et al*, 2014; Huttlin *et al*, 2015).

To further benchmark our PPI-network, we investigated whether it was enriched for reported protein complexes. With only 19 SLCs covered, SLCs and their complex associations are underrepresented within CORUM (Tsitsiridis *et al*, 2023), a reference database of protein complexes, in comparison to other TM or soluble proteins (**Fig. EV1E**). For this we assumed full connectivity between all interactors of each SLC and of all complex subunits reported in CORUM. We intersected the CORUM derived PPI-network against 10,000 permuted networks with the same topology and composition as the SLC- interactome (see methods). CORUM interaction pairs were significantly enriched within the SLC- interactome compared to permutated networks (**Fig. 2H**).

### Orthogonal evidence of SLC-protein interactions by *in silico* structural modelling

Recent progress in deep-learning methods to predict structures of experimentally determined protein interactions can lead to high-confidence structural models of protein complexes (Burke *et al*, 2023). However, the success rate for the long C- and N-terminal tails of TM-proteins is limited, as these tails often contain intrinsically disorder regions (IDR) with protein interactions mostly mediated by small linear motifs (SLiMs) (Morris *et al*, 2021). Despite these limitations, we used AlphaFold-Multimer for structural prediction of PPIs (Evans *et al*, 2021). For the *in silico* validation, we decided to start from well-studied SLC-protein interactions and expand to novel interactions of SLCs with the same proteins.

First, we focused on heterodimeric complexes of monocarboxylate transporters (MCTs). SLC16A- family members were reported to form stable associations with small single-pass transmembrane chaperone proteins Basigin (BSG) and Embigin (EMB) (Bosshart *et al*, 2021). It was further reported that the SLC16A11 interaction with BSG was dysregulated in Type 2 diabetes variants (Rusu *et al*, 2017). In our data set, BSG was detected in 93.33% of all AP-MS samples and scored for eleven SLCs. We found interactions with several SLC16A-family members and additionally with SLCO1B1, SLCO1B3, MFSD8 and SLC19A1. Compared to BSG, EMB was only detected in ∼20% of all our AP-MS experiments (**Fig. 2I**). We modelled the structure of the SLC-chaperone interactions found in our SLC-interactome (**Fig. 2J, Appendix Fig. S3A-D**). For 13 out of the 15 heterodimers we retrieved high-confidence structures (pDockQ > 0.5), among them SLC16A1 in complex with both chaperones as previously reported (Halestrap, 2013), as well as SLC16A5 and SLC16A11 in complex with BSG, for which no structure was reported. The amino acid transporter heavy chain (SLC3A2), is another well-known subunit of heterodimeric SLC-complexes formed with multiple members of the cationic amino acid transporters (CATs) and glycoprotein-associated amino acid transporters (gpaATs) of the SLC7A-family (Rodriguez *et al*, 2021; Lee *et al*, 2022; Oda *et al*, 2020). SLC3A2 was detected in 87% of all SLC AP-MS experiments and scored as interaction partner for a total of seven SLCs, of which six are members of the SLC7A-family (**Fig. EV1F**). Our data set recapitulates almost all known SLC3A2 interactions, missing only the interaction with SLC7A9, for which the SLC3A2 signal was slightly lower compared to the scored interactions and was thus classified as background. We further modelled the complexes and obtained high-confidence structures for all SLC3A2 interactions (**Fig. EV1G**). Comparing predicted chaperone complexes against a set of non-reported chaperone-SLC interactions, randomly sampled from the lowest 25% quantile of SLC3A2 abundance quantified within the SLC-interactome, showed that the scored interactions led to significantly higher scoring structures (unpaired student t-test, p-value=0.0009177, **Fig. EV1G right side, Appendix Fig. S4A,B**).

A third example of heterodimeric complexes formed by several family members was the calcium-binding protein CHP1 and the Na+/H+ exchanger transporters SLC9A1 and SLC9A3 of the SLC9A-family. CHP1 was reported to stabilize their plasma membrane localization and to increase pH sensitivity (Dong *et al*, 2021). We found CHP1 to interact with even more SLC9A-family members as well as SLC6A- and SLC7A-family members (**Fig. EV1H**). CHP1 was detected at significant levels for SLC9A2– SLC9A4 and with lower abundance for SLC9A5–SLC9A9. The affinity purifications of SLC9A1 did not yield sufficient signal and were thus excluded from the SLC-interactome study. Structural prediction of these PPIs resulted in high-confidence structures for all CHP1-SLC9A-family heterodimers and for four other SLCs with a medium confidence structure (**Fig. EV1I**). Also, the obtained structural models for the identified SLC-CHP1 interactions were of significantly higher quality compared to a random set of SLC-CHP1 interactions classified as background (unpaired student t-test, p-value=0.03642, **Fig. EV1I** right side, **Appendix Fig. S4C,D**).

In summary, the benchmarking effort highlighted that the SLC-interactome recapitulated well-characterized interactions and recovered thousands of novel interactions. The recovery rate of already reported interactions was higher compared to other large-scale studies. Prediction of heterodimers enabled the *in silico* validation of several known as well as unreported SLC-containing complexes. Finally, this SLC interactome combined with machine learning algorithms is suitable for predicting the structure of interacting surfaces and for the generation of thousands of three-dimensional models, which are useful for interpreting SLC genetic variants associated with disease.

### Clustering of SLC-interaction profiles reveals functionally relevant interaction networks

Faced with the challenge of visualizing the interactome in a manner that is reader-friendly and biologically meaningful, we abstained from a hairball network representation. Clustering of interactor relations is regularly used to deconvolute high-density PPI-networks, obtained by AP-MS (Buljan *et al*, 2020; Uliana *et al*, 2023) or *in vivo* proximity interaction mapping (Go *et al*, 2021; Salokas *et al*, 2022), into protein complexes. In addition to this approach, we grouped the SLCs based on their PPI-network similarity to derive common properties such as subcellular location, protein complexes and family-wide interactions.

For the latter approach, we performed hierarchical clustering based on the overlap of interaction partners, resulting in a dendrogram of 396 SLCs in 38 clusters (**Fig. 3A)**. The number of clusters was chosen using a local peak in the mean silhouette width by estimating a cluster number from k=2 to k=110 **(Appendix Fig. S5**). The clusters included on average 10 SLCs (min: 2, max: 78) and 319 interactions (min: 2, max: 1,185).

**Figure 3.**
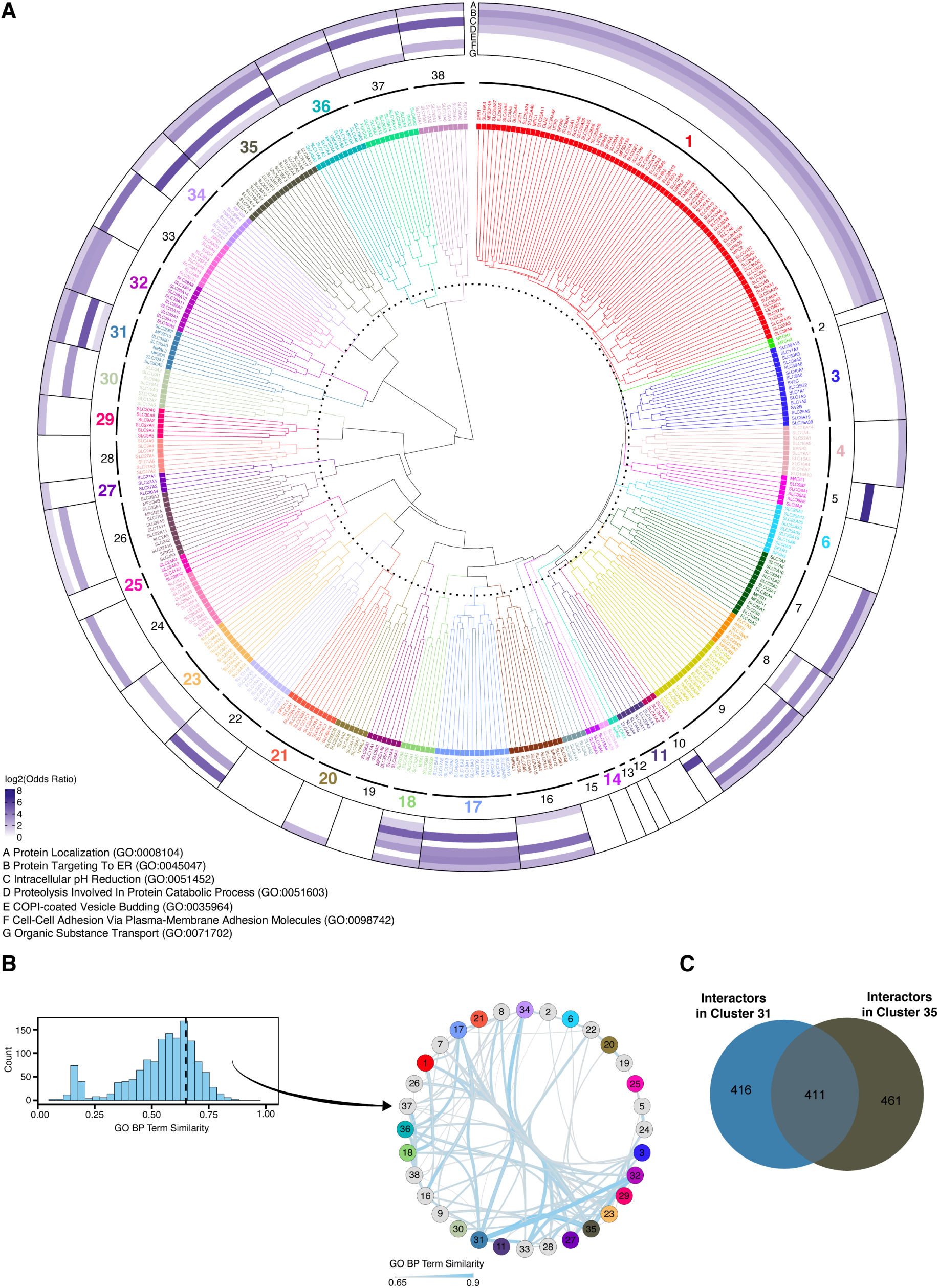
Clustering SLCs by their interactome similarity. **(A)** Dendrogram of hierarchical clustering based on the Jaccard similarity matrix derived from the interactome profiles of 396 SLCs. The heatmap displays the log ratio of the top-level biological process enriched pathways. Clusters with a significantly enriched SLC functional property (Fisher’s test p < 0.2) are shown in bold and the respective cluster colour. Outer ring shows representative parental GO terms significantly enriched within the cluster (p-value < 0.01). **(B)** Distribution of GO semantic similarities between SLC similarity clusters. A similarity threshold of 0.65 (dashed black line) between the clusters was chosen to filter the data. **(C)** Overlap of protein interactors identified among the SLCs from cluster 31 and cluster 35.

For a coarse overview of biological processes associated with the interactome profiles of each cluster, we performed GO biological processing terms enrichment analysis. To reduce complexity and graphically represent functional similarities between different SLCs, we filtered for frequent terms and then performed hierarchical clustering in combination with semantic similarity and reduction analysis to determine important parental terms. We obtained seven representative parental terms to annotate the PPI-network dendrogram (**Fig. 3A outer ring**, **Appendix Fig. S6B,C** and methods).

Most of the parental terms were enriched for clusters 17, 18, 31 and 35, indicating that they contain SLCs with large, complex interactomes. Other clusters were only enriched for specific terms; for example, clusters 5 and 34 showed a strong enrichment for COPI-coated Vesicle Budding. The parental terms proteostasis and protein subcellular localization and trafficking were more often found to be significant. Some clusters lack enrichment for any parental term, as we removed child terms to simplify visualization. A heatmap of all enriched terms is shown in **Appendix Fig. S6B**.

To study the relationship between the clusters, we used all enriched terms (5,675 terms, p-value < 0.01) and derived a similarity network based on GO semantic similarities (**Fig. 3B**). The network retained 32 of the 38 clusters, and clusters 31, 17 and 35 were the most connected. Comparing clusters 31 and 35 on SLC level showed that both contained SLC35 family members. After overlapping the PPI-networks of the two clusters, we found that roughly a third (411) of all interaction partners are present in both clusters (**Fig. 3C**). Among the interactors found in at least 50% of all SLCs in these two clusters were RABL3 and ERMP1, associated with trafficking/signalling and an ER residing endopeptidase.

To further compare the SLCs, we tested by enrichment analysis whether SLCs were grouped according to functional properties such as fold, localization, substrate class or family membership (methods for details; SLC annotation from Goldmann et al, accompanying manuscript). For 20 of the 38 clusters (53%) we retrieved at least one significantly enriched functional property (Fisher’s test p < 0.2) (**Fig. EV3A**) and in total, we found 44 significantly enriched SLC properties that were grouped into five distinct categories: coupled ion, family, fold, location, and substrate (**Fig. EV3B**). SLC families were overrepresented in several clusters (e.g. SLC4, SLC6, SLC27 and SLC39), indicating that these SLCs share more interactions within their family compared to other SLCs in the SLC-interactome (**Fig. EV3A**). For cluster 30, multiple properties were enriched: coupled-ion, substrate, family and type of fold (**Fig. EV3C**). The cluster contained eight members from SLC12A and SLC6A families, annotated as chloride dependent transporters (Meixner *et al*, 2020), and their combined PPI-network covered a subset of 231 interactions (**Fig. EV3D**). Enrichment analysis of proteins within the cluster identified cell volume regulation, chloride and ion homeostasis, protein folding and chaperone mediated protein complex folding as significant terms (**Fig EV3E**), thus linking the interaction partners recovered in cluster 30 to the transported substrates of its members (Song *et al*, 2002; Syringas *et al*, 2000). 21 out of the 25 most connected interactors in the cluster specific PPI-network are reported as physically interconnected (STRING score > 0.4, filtered for physical interactions; **Fig. EV3G**). Several of them were folding chaperones, including two HSP90 subunits and four HSP70 isoforms (**Fig. EV3F**). It was previously found that the immature, ER-resident form of SLC12A1 interacted with HSP70 (Bakhos-Douaihy *et al*, 2021) as well as other family members relying on chaperoning for trafficking (Rogala-Koziarska *et al*, 2019). These shared chaperone interactions likely represent one layer of ER quality control during the folding process of SLC12 and SLC6 members, possibly because SLC12 family members are large proteins that require specialized stabilization and folding of the LeuT fold.

As HSP70 and HSP90 multichaperones were quantified in many of the SLCs, and particularly high in SLC12A-family members, we next investigated if there is a general dependence of finding this interaction and the SLC-protein length. Comparing the protein length of all SLCs interacting with HSP90AA1 with SLCs for which HSP90AA1 was only recovered in the background showed that the median protein length of SLCs interacting with the chaperone was significantly longer (1,084 versus 533 amino acids; student t-test p-value=1.285 x 10^-8^, **Fig. EV3H**). Next, we tested if the abundance of chaperones correlated with the SLC tail length (methods for details). This revealed that 77% of the chaperone showed a weak to moderate relationship (**Fig. EV3I**).

To reveal complexes covered by multiple SLCs, we clustered interaction partners across the SLC- interactome. We pre-filtered all interaction partners by their connectivity, resulting in a subnetwork of 2,835 interactors from 371 SLCs, and derived a correlation-based distance matrix. We performed unsupervised hierarchical clustering (Ward D2, silhouette plot **Appendix Fig. S7A**) resulting in 207 clusters (**Appendix Fig. S7B**). Next, we performed GSEA for GO biological process terms on the cluster members and found significant terms for 124 (60%) clusters (adjusted p-value ≤ 0.01) (**Appendix Fig. S7C**). The top five terms were in line with the coarse overview of biological processes associated with SLC-clusters used in **Fig. 3** (**Appendix Fig. S7D**). To test if interactors of the same cluster form a complex, we intersected CORUM complexes with the interactor clusters, and filtered for completeness of complexes using a 40% complex completeness threshold (**Appendix Fig. S7E**). For 73% (151) of the clusters, we found a total of 796 CORUM complexes (680 unique CORUM complexes), with the Rag-Ragulator complex being the most complete (Rebsamen *et al*, 2015). In addition, we found several complexes, associated with trafficking, including multiple small LIN-complexes (**Appendix Fig. S7F**). The LIN-complex will be discussed in more detail in the trafficking result section.

In summary, similarity clustering combined with functional annotation of SLC-cluster specific networks allowed us to relate SLCs to each other on a functional level. Dissecting one of them lead to the identification of a set of chaperones for SLC12A and SLC6A family members, likely involved in their protein quality control. The correlation analysis of interactors identified several protein complexes, despite the sparseness of the SLC-interactome, among them the LIN-complex.

### Proteostatic regulation of SLCs

Any functional validation strategy of the SLC interactome, on such a broad spectrum of transporters, localised on different subcellular membranes and likely to transport different substrates, risks of being exemplary and anecdotal in nature. From the pharmacological perspective inspired by the mission of RESOLUTE, modulation of the proteostatic regulation of transporters may restore function to poorly folded or mislocalised gene variants associated with human disease. We therefore considered a strategy allowing to assess new interactors for their ability of affecting the localisation and protein levels of transporters. Indeed, GO term enrichment analysis for biological processes of all detected interactions in the SLC interactome identified these biological themes recurring among the top terms (**Fig. 3A, Appendix Fig. S6A,B**).

We devised a validation workflow that assessed protein localization and degradation. We selected 65 SLCs with interactions involved in these processes and expressed them with a GFP tag from an inducible promoter (**Fig. 4A**). To measure protein stability by flow cytometry (**Fig. 4B**), we added an RFP reference protein linked by an internal ribosome entry site to the expression cassette (Yen *et al*, 2008). To measure changes in subcellular localization of the SLC by microscopy, we further modified the reference-RFP to localize either to 1) the entire cell body; 2) the plasma membrane; 3) mitochondria; or 4) lysosomes, depending on the SLC localization (Goldmann et al, accompanying manuscript; see also **Fig. 6, Table EV3**). By using RNA-based interference (loss-of-function) or cDNA overexpression (gain-of-function) of the interactor we tested effects on localization or abundance of the cognate SLC.

**Figure 4.**
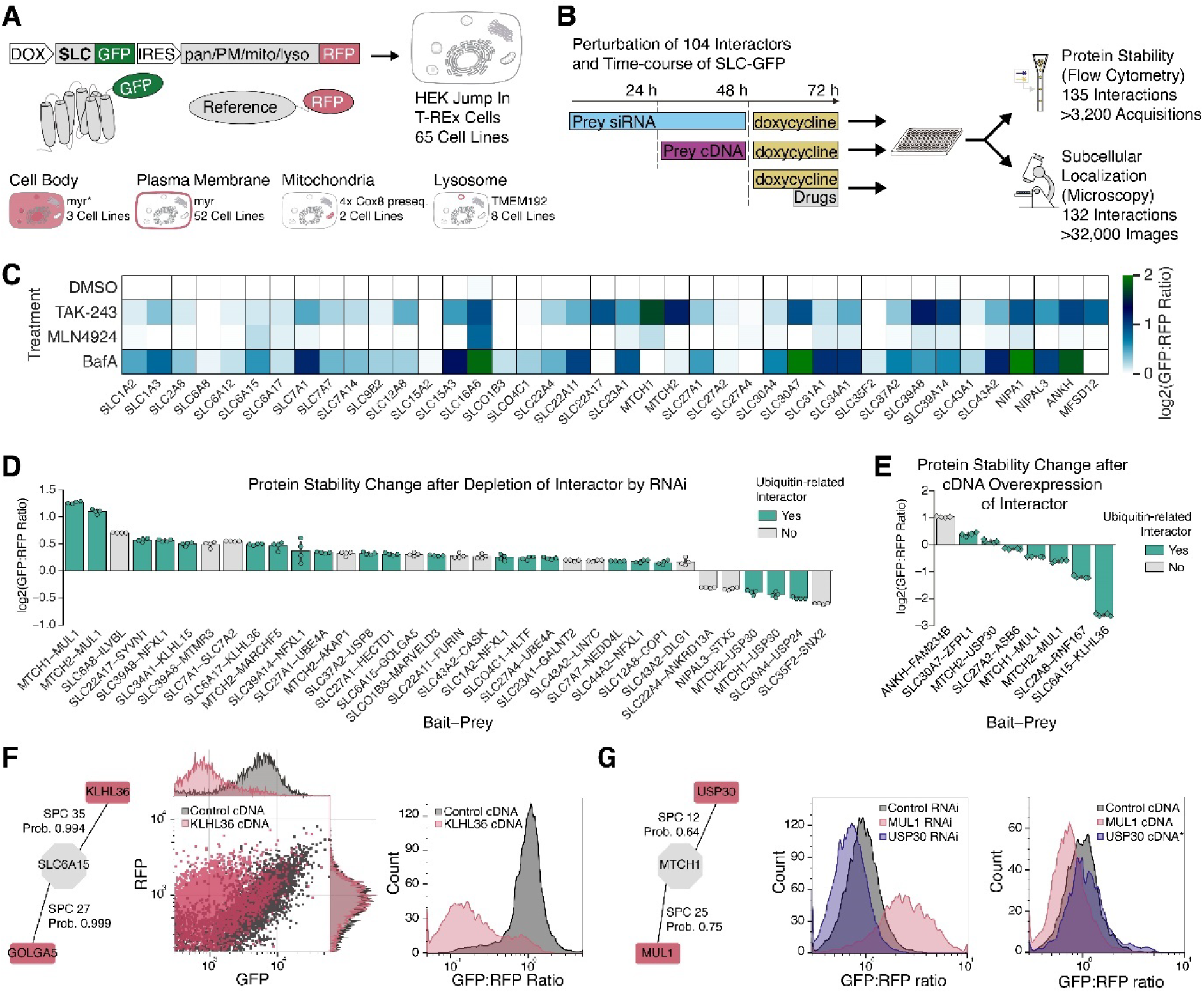
Proteostatic regulation of SLCs. **(A)** Generation of cell lines conditionally expressing GFP-tagged SLCs together with an RFP reporter modified with membrane-specific tags or proteins that allow localization to biologically relevant membranes. **(B)** Protein stability and subcellular localization assay to validate SLC interactions. Selected SLCs were conditionally expressed and monitored using flow cytometry and microscopy. GFP:RFP ratio and GFP localization and intensity were measured after perturbation of interactor abundance or drug treatments. **(C)** SLC protein stability after drug treatment as measured by flow cytometry showed distinct susceptibility to degradation pathways. Heatmap of log2 transformed GFP:RFP ratios measured after 24 h of induction and drug treatment in the last 6 hours (normalized to DMSO control; n=3 wells per sample). **(D)** Results for 37 SLC-protein interactions with significant changes to GFP:RFP ratio values out of 135 tested interactions by RNAi, showing mean values ± 95% confidence intervals (n=4 wells per condition, p-value < 0.01, independent t test; GFP:RFP and GFP:GFP ratios compared to control treatment changed to ≤90% or ≥110%; see Methods for details). **(E)** Results for 9 SLC-protein interactions after interactor cDNA overexpression out of 18 tested interactions analysed as in (D). **(F)** KLHL36 and GOLGA5 were strongly enriched in SLC6A15 purifications compared to other SLCs or GFP (n=2 biologically independent replicates). In flow cytometry experiments, SLC6A15-GFP was destabilized by overexpression of the ubiquitin ligase adapter KLHL36 (n > 4,600 events per sample). **(G)** MTCH1 was destabilized by E3 ligase MUL1 and stabilized by deubiquitinase USP30. In flow cytometry experiments, MTCH1-GFP protein stability directly correlated with depletion by RNAi or cDNA overexpression of its interactors and their biological functions in the ubiquitin-dependent degradation pathway (n > 2,500 events per sample). Asterisk indicates an effect that was below the set threshold of 10% change in GFP:RFP ratio. See also **Fig. EV4**.

To characterize the cell lines and better understand the effects of protein degradation on the SLCs, we treated 40 of these cell lines with inhibitors of the ubiquitin pathway (TAK-243; (Hyer *et al*, 2018)), the neddylation pathway (MLN4924; (Soucy *et al*, 2009)), or lysosomal acidification (Bafilomycin A1), and measured changes to the GFP:RFP ratio relative to DMSO treatment (**Fig. 4C**). SLC levels were either unchanged or stabilized by these inhibitors. Applying a threshold of 10% increase in protein levels, only SLC16A6 was stabilized by all three inhibitors, whereas most SLCs were specifically stabilized by either inhibiting ubiquitination, lysosome acidification, or both (**Fig. EV4A**). Five SLCs showed no significant stabilization after any of the selected treatments, suggesting that they are not regulated by these degradation pathways in HEK 293 cells (**Fig. EV4A**).

Next, we measured the protein stability of 65 GFP-tagged SLCs after 134 treatments with RNAi and 18 with transient cDNA overexpression, resulting in 152 conditions that we classified into changed or unchanged protein levels. We set a threshold of 10% change in mean GFP:RFP ratio and overall GFP levels compared to control treated samples at a p value < 0.01. We also excluded measurements if relative RFP levels changed 50% or more of relative GFP levels in the same direction (**Dataset EV2**). We thus found 37 interactions that showed robust changes in GFP:RFP ratios after RNAi-mediated depletion (**Fig. 4D**) and 9 interactions after cDNA-mediated overexpression (**Fig. 4E**). These observations include 29 protein interactions that are functionally linked to ubiquitination, where we found that depletion of E3 ligases and adaptor proteins increased SLC levels and depletion of DUB proteins decreased levels in all but three cases. Notably, eight of these interactions were previously reported in large scale studies (of which four PPIs were identified in AP-MS; **Dataset EV2**). To our knowledge, none had been functionally assessed before.

We also tested pharmacological inhibition of degradation in addition to depleting or overexpressing the SLC interactor in HEK 293 cells (**Appendix Fig. S8**). The increase in protein levels observed after RNAi-mediated depletion was not further increased in some cases, suggesting that the main degrader of a specific SLC had already been eliminated (e.g. SLC2A8–RNF167) or that the interactor was not directly linked to these degradation pathways (e.g. SLC6A8–ILVBL; SLC6A15–GOLGA5). In other cases, SLC levels were additionally increased by inhibiting ubiquitination or lysosome acidification (e.g. MTCH2–MUL1; SLC39A8–NFXL1), indicating either incomplete depletion, additional degraders or other stabilization mechanisms.

SLC6A15 was stabilized after treatment with Bafilomycin A1 and MLN4924 (**Fig. EV4A**) and showed the strongest decrease in stability upon overexpression of its interactor KLHL36 with a reduction of the SLC6A15-GFP to RFP ratio by 6-fold **(Fig. 4E,F**). KLHL36 is a potential cullin-based E3 ligase adaptor, and uniquely interacts with SLC6A15 and SLC6A17 in the SLC-interactome data set (**Fig. EV4B**). Addition of any of the three inhibitors six hours preceding analysis partly stabilized the GFP signal, indicating that SLC6A15 protein stability is regulated by a KLHL36-associated cullin-based RING E3 ligase and the autophagy-lysosomal pathway (**Fig. EV4H**). Using an antibody against endogenous

SLC6A15 confirmed the degradation by KLHL36 overexpression (**Fig. EV4C**). These results suggested a protein degradation mechanism that specifically employs a Kelch domain protein, cullin-RING-ligases, and lysosomes. Other SLCs may also involve a combined degradation mechanism since close to half of the SLCs tested were stabilized by both Bafilomycin A1 and TAK-243 (**Fig. EV4A**).

The mitochondrial carrier protein MTCH1 was the most strongly stabilized SLC following pharmacological inhibition of ubiquitination (**Fig. 4C**). When depleting the interactors mitochondrial ubiquitin ligase activator of NFKB1 (MUL1) and Ubiquitin carboxyl-terminal hydrolase 30 (USP30) by RNAi, MTCH1-GFP was stabilized 2.4-fold and destabilized by 1.4-fold, respectively (**Fig. 4G**). Transient overexpression resulted in the opposite effect, with the E3 ligase reducing and the deubiquitinase increasing MTCH1-GFP levels (**Fig. 4G**). We obtained similar results for MTCH2-GFP which also interacted with MUL1 and USP30; however, we found that MTCH2 stability was also regulated by another E3 ligase, MARCHF5 (**Fig. EV4D, Fig. 4D,E**). To investigate this further, the stabilization of MTCH2 was also assessed in a HAP1 cell model where endogenous MTCH2 was tagged at its N-terminus with HA-dTAG (HA-FKBP12^F36V^). After depletion of MARCHF5 or MUL1, endogenous MTCH2 stabilization was observed in bulk via Western blotting (**Fig. EV4E**) and specifically at mitochondria via immunofluorescence (**Fig. EV4F**).

These results highlight the complexity of SLC proteostasis, involving many different protein classes across a wide range of SLC families and suggesting a prominent role of both the proteasome and the lysosome in SLC degradation.

### Destabilization of monocarboxylate transporter SLC16A6 by a phospho-degron

The strong increase of SLC16A6 (MCT7, Monocarboxylate transporter 7) protein levels caused by the neddylation inhibitor MLN4924 suggested involvement of the SCF (SKP1-CUL1-F-box) ubiquitin ligase known to be activated by neddylation (**Fig. 4B**). The interactome of SLC16A6 contained the CUL1 subunit and its two F-box adaptor proteins BTRC and FBXW11 (Frescas & Pagano, 2008) (**Fig. 5A**), which were unique to SLC16A6 (**Fig. EV5A, B**). The essential SCF subunit SKP1 was found in the SLC16A6 purification, but since it was found in most SLC purifications and is considered a common contaminant (Mellacheruvu *et al*, 2013), it was removed post-acquisition (**Fig. EV5B**). Enrichment analysis of SLC16A6 interactors for GO biological processes resulted in a significant enrichment for the SCF-dependent proteasomal ubiquitin-dependent process term (**Fig. EV5C).**

**Figure 5.**
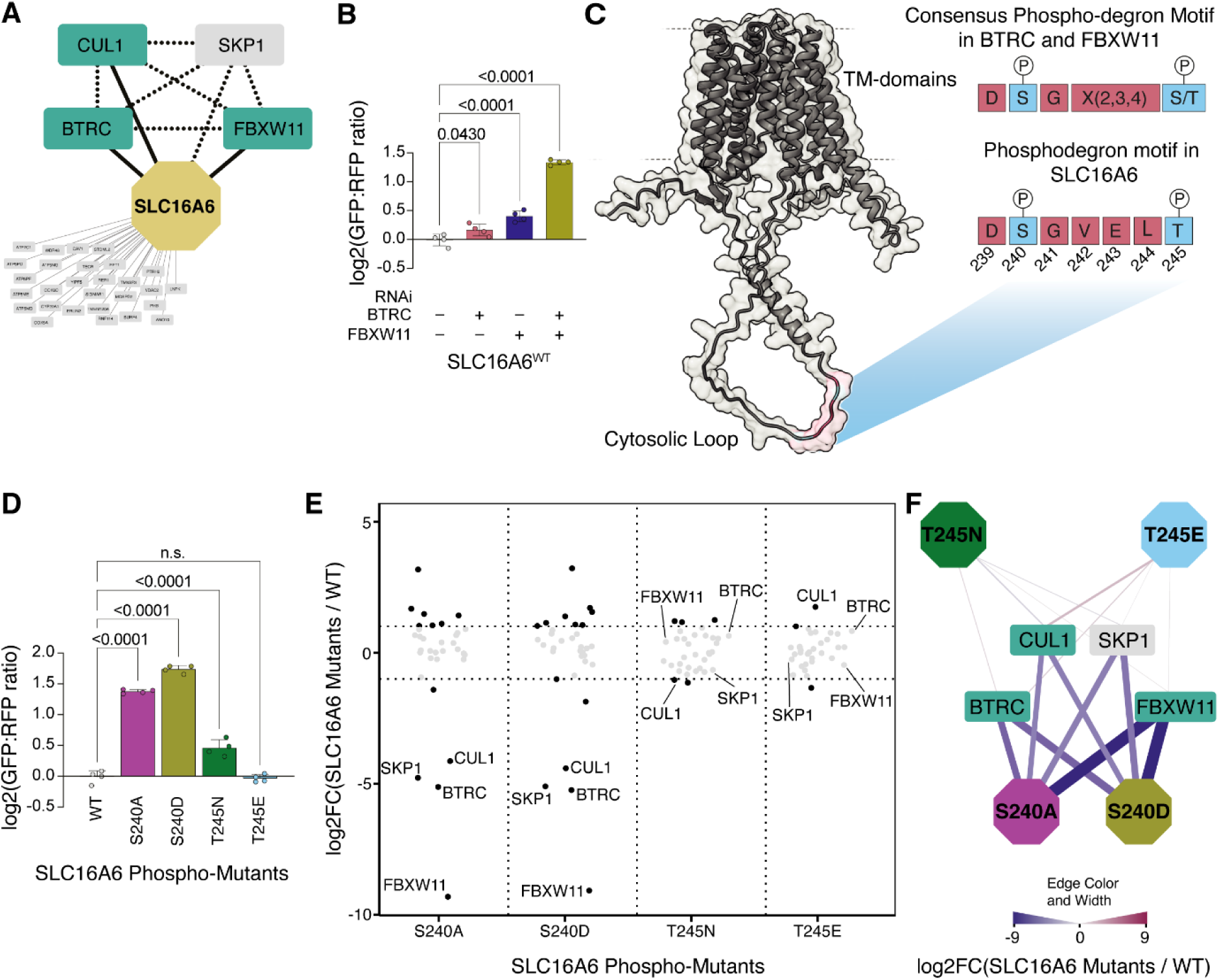
SLC16A6 stability is regulated by a phospho-degron. **(A)** SLC16A6 interacts with subunits of E3 ubiquitin ligase SCF. SLC16A6 coloured in beige, interactors in green. SKP1 (grey) was quantified but not scored and supplemented to the network. Novel edges depicted by solid black lines and edges obtained from literature in black dotted lines. **(B)** Protein stability of SLC16A6 after RNAi treatment of adaptor proteins showing strong stabilization after co-depletion of BTRC and FBXW11 (mean ± SD; p-values were calculated using one-way ANOVA and Dunnett test; n=4 replicates). **(C)** Alphafold predicted structure of SLC16A6 with highlighted cytosolic loop containing the phospho-degron. The βTrCP consensus motif and SLC16A6 phospho-degron motif are visualized on the right side. **(D)** Protein stability assay of SLC16A6 phospho-mutants compared to SLC16A6^WT^ (mean ± SD; p-values were calculated using one-way ANOVA and Dunnett test; n=4 replicates). **(E)** Quantitative comparison of SCF complex subunits across SLC16A6 phospho-mutants (x-axis) against SLC16A6^WT^. The log2FC (y-axis) were derived for the 31 interaction partners against their abundance in SLC16A6^WT^. Dashed black line indicates a ±1 log2FC threshold. Labelled dots represent SCF-complex members (n=4). **(F)** PPI-Network of SLC16A6 phospho-mutant interactions with SCF complex subunits. The colour and thickness of edges indicate the mean log2FC against SLC16A6^WT^. For all AP-MS data shown in the figure panels, n=2 biologically independent replicates with n=2 technical injections were used.

Depletion of FBXW11 increased the median levels of GFP-SLC16A6 by 8.5%, below our set threshold for robust changes (**Fig. EV4C**). Because BTRC and FBXW11 (also known as β-TrCP and β-TrCP2, respectively) are paralogs thought to have redundant roles (Frescas & Pagano, 2008), we analysed SLC16A6 protein stability after single and combined RNAi treatment of the two F-box proteins (**Fig. 5B**). Combined treatment with RNAi against BTRC and FBXW11 stabilized SLC16A6 by 1.5-fold compared to control RNAi treatment (**Fig. 5B**). Additional treatment with MLN4924 did not significantly increase protein levels (**Fig. EV5D**), indicating that distribution of the SCF^β-TRCP^ complex was mainly responsible for the observed stabilization of GFP-SLC16A6.

Protein degradation via E3 ligases often occurs via specific linear sequence motifs or degrons (Sherpa *et al*, 2022). Substrates of BTRC and FBXW11 share the consensus sequence DSGX(2,3,4)S/T in which the serine residues are phosphorylated to allow binding and subsequent ubiquitination and degradation **(Fig. EV5E)** (Cardozo & Pagano, 2004; Low *et al*, 2014); interestingly, SLC16A6 contains a DSGVELT motif at positions 239–245 (**Fig. 5C**). Experimental structures of SLC16A6 are not yet available, but a structural model by AlphaFold2 mapped this motif to a cytosolic loop of the transmembrane protein where it could potentially be phosphorylated and recognized by the SCF^βTrCP^ complex (**Fig. 5C**).

To investigate whether the motif in SLC16A6 is a phosphorylated degron we individually mutated serine 240 to alanine (S240A) and threonine 245 to asparagine (T245N). We also generated a serine to aspartate (S240D) and a threonine to glutamate (T245E) mutant to potentially mimic phosphorylated residues at these positions and elicit increased degradation. The non-phosphorylatable S240A and T245N mutations increased SLC16A6 levels assessed by flow cytometry, suggesting that the motif targets SLC16A6 for degradation (**Fig. 5D**). The S240D mutant was the most stable, suggesting that the aspartate substitution cannot substitute for phosphorylated serine. The T245E mutant was degraded as efficiently as unmodified GFP-SLC16A6 (**Fig. 5D**). Assessing SLC16A6 localization by live cell microscopy showed accumulation of the S240A and S240D mutants at the plasma membrane (marked by expression of modified RFP), in line with the flow cytometry results (**Fig. EV5F**). While the T245N mutant only showed slight stabilization, the T245E mutant was even less abundant at the plasma membrane than wild type SLC16A6, suggesting either better recognition by the SCF and improved degradation, or changes in subcellular SLC16A6 distribution (**Fig. EV5F**).

We depleted BTRC and FBXW11 in the mutant cell lines to ensure that no other degrons were operational (**Fig. EV5G**). Simultaneous depletion of the two F-box proteins by RNAi did not further stabilize the S240A and S240D mutants, suggesting that serine 240 is the crucial residue determining SLC16A6 stability. The T245N and T245E mutants and wild type SLC16A6 were stabilized by BTRC and FBXW11 depletion, suggesting that all three proteins encode functional degrons. To investigate whether degron mutations decrease the interaction with SCF proteins, we performed AP-MS on the SH-tagged SLC16A6 mutants. BTRC, CUL1 and FBXW11 were strongly depleted or absent from serine mutants, while the T245E mutant showed increased CUL1 and equal levels of the other SCF subunits as wild type SLC16A6 (**Fig. 5E, 5F, Fig. EV5H, Dataset EV3**). The differential interactome data further emphasized the requirement for SLC16A6 serine 240 and its phosphorylation to ensure efficient recognition of the resulting degron and direct binding by BTRC and FBXW11.

### Proteins affecting trafficking and subcellular localization of SLCs

To reach their site of action, transmembrane proteins interact with a diverse set of proteins and traverse several subcellular compartments. To examine if this is reflected in the SLC-interactome, we performed gene set enrichment analysis of cellular compartment ontology terms and summarized the terms into ten subcellular environments (p < 0.05; **Fig. EV6A**). More than half of SLCs interacted with proteins associated with the ER, with vesicles, and with the plasma membrane where the majority of SLCs localize (Meixner et al. 2020). On average, each SLC interactome was associated with 4.5 out of 10 subcellular compartments, independent of reported intracellular or plasma membrane localization (**Fig. EV6B**, **EV6C**), suggesting that AP-MS purification represented a composite of several SLC life cycle locations. Clustering the SLCs based on their respective enriched compartments revealed, among others, subsets of mitochondrial (cluster 20) and lysosomal (cluster 24) SLC interactomes (**Fig. EV6D**).

To understand their role in trafficking of SLCs, we depleted individual interactors by RNAi and analysed the subcellular localization of GFP-tagged SLCs in live cells, with good correlation between our cytometry and microscopy assays **(Fig. 4B, Appendix Fig. S9**). Using a co-expressed reference RFP we calculated the overlap between SLC-GFP and reference signals and measured changes in GFP intensity at RFP positive and negative regions. Out of 132 unique interactions analysed, 73 interactions showed no changes to the SLC-GFP signal after depletion of the interactor. For 59 interactions, corresponding to 36 SLCs, we observed significant changes to the GFP levels or subcellular GFP signal distribution (**Dataset EV2**).

Among them, thirteen SLCs localized to a single subcellular compartment or were covered by the cell body reference RFP. Interactor depletion for 25 SLC–interactor combinations led to SLC-GFP intensity changes at this location (**Fig. 6A**). This is exemplified by SLC22A4 and its interaction with the trafficking regulator YIPF3 that when depleted, significantly reduced the SLC22A4-GFP signal at the plasma membrane (**Fig. 6A**). In another case, we measured SLC1A2 signal at the plasma membrane after depletion of three of its interactors. GFP intensity decreased after GDPD1 and NEK6 RNAi (**Fig. 6B**). When we used an impedance-based assay of cell morphology (Sijben *et al*, 2022) to monitor transporter activity of SLC1A2, SLC1A3 and SLC22A3 (**Appendix Fig. S10**), we found that GDPD1 RNAi significantly decreased glutamate uptake in HEK 293 Jump In T-REx cells overexpressing SLC1A2 (p-value = 0.0121) (**Fig. 6C**).

**Figure 6.**
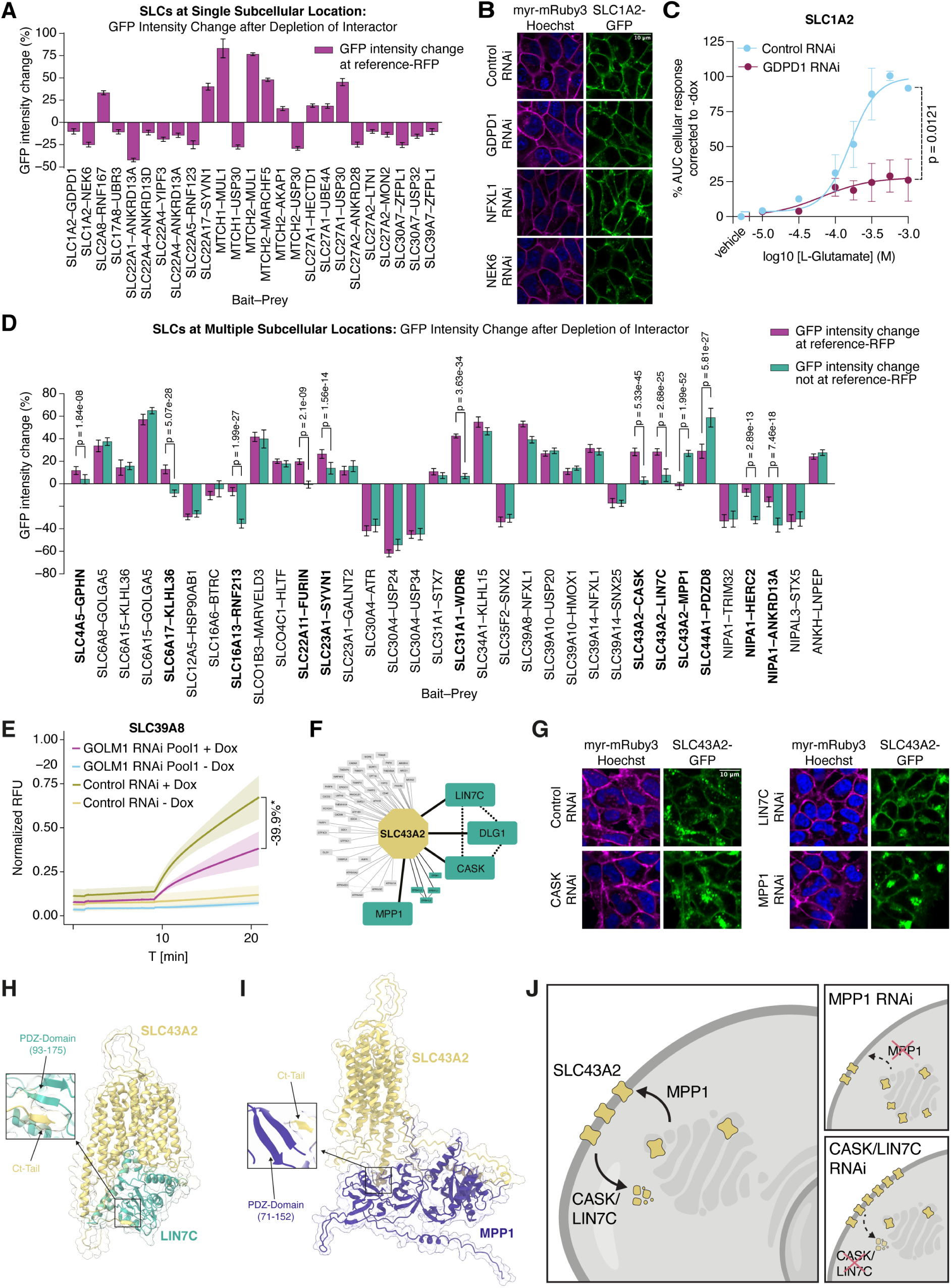
Proteins affecting trafficking and subcellular localization of SLCs. **(A)** Relative GFP intensity changes at reference regions after interactor RNAi compared to control RNAi are plotted as a percentage (n=160 images per condition; error bars denote 95% confidence intervals). The graph includes SLCs residing at single subcellular locations and SLC30A7 (since vesicles and Golgi locations are both covered by expression of cell body RFP). GFP intensity changes at the respective reference signal were thresholded at 10% increase or decrease over control treatment with p-value < 0.01 (independent t test). Subcellular compartment location of SLC-GFP and reference RFP are indicated in **Dataset EV2**. **(B)** Representative images of plasma membrane localized SLC1A2-GFP after depletion of 3 interactors (scale bar, 10 um). GFP signal intensity was quantified using myr-mRuby3 to identify plasma membrane pixels. **(C)** Quantification of cellular impedance of HEK 293 SLC1A2-SH cells treated with increasing concentrations of glutamate after GDPD1 RNAi. Data were corrected against samples without doxycycline due to significant L-Glutamine transport by endogenous SLC1A3 in HEK 293 cells and are shown as background corrected mean area under the curve (AUC) quantification relative to control RNAi ±SEM. p-value was calculated using unpaired t test. **(D)** Relative GFP intensity changes at the reference region (purple) and outside the reference region (teal) for 34 SLC-interactor RNAi combinations for SLC-GFP signals covering more than one subcellular compartment (as indicated in **Dataset EV2;** n=160 images per condition; error bars denote 95% confidence intervals). The graph includes interactions with GFP intensity changes exceeding 10% at the reference region or 25% outside the reference region with p-values < 0.01 compared to control RNAi (independent t test). For 13 interactions highlighted in bold, GFP intensity changes at reference and non-reference regions differed significantly (p-values as indicated, independent t test with Benjamini-Hochberg correction for multiple tests; relative change at least 33.3% of absolute higher value), indicating location-specific changes. GFP intensity changes for other interactions were similar for reference and non-reference regions, indicating general changes in SLC abundance after interactor depletion. **(E)** Relative fluorescence unit (RFU) curve to measure cadmium uptake of SLC39A8 after RNAi against GOLM1. Traces represent mean min-max normalized RFU values across replicates ±SD (curve shades; n=3 biologically independent replicates with 8 technical replicates). Results are compared to control RNAi using mean AUC calculated from normalized RFU traces considering all replicates (**Appendix Fig. S11**). **(F)** SLC43A2 interactions with trafficking proteins CASK, DLG1, LIN7C and MPP1 (teal; n=2 biological replicates with n=2 technical replicates). Novel edges are depicted by solid lines and edges supplemented from BioGRID are shown with dotted line. **(G)** Representative images of plasma membrane and vesicle localized SLC43A2 after depletion of 3 interactors (scale bar, 10 µm). Depletion of CASK or LIN7C increased plasma membrane associated GFP signal whereas RNAi of MPP1 increased the intensity and relative distribution of SLC43A2-GFP away from the plasma membrane (quantified in panel D). **(H)** AlphaFold2 model of SLC43A2– LIN7C complex with section of the interaction of the LIN7C PDZ domain and the intracellular C-terminal tail of SLC43A2 (Weighted Score 0.57). **(I)** AlphaFold2 model of SLC43A2–MPP1 interaction between the MPP1 PDZ domain and the intracellular C-terminal tail of SLC43A2 (Weighted Score 0.51). **(J)** Schematic of SLC43A2 trafficking regulation.

For 22/34 interactions of 23 SLCs localized to multiple compartments, SLC-GFP levels changed indiscriminately at all subcellular locations (**Fig. 6D**). For instance, plasma membrane- and ER-associated GFP signals for SLC39A8 increased upon depletion of NFXL1. We further tested if the increase affected SLC39A8 transport in a cadmium uptake assay. While NFXL1 RNAi did not change cadmium uptake, depletion of the interactors GOLM1 and, to lesser extent, sorting nexin 25 (SNX25) conferred a significant reduction in fluorescent signal, corresponding to reduced cadmium uptake (**Fig. 6E**, **Appendix Fig. S11**). Since GOLM1 and SNX25 did not measurably change SLC39A8 localization or protein levels, they likely regulate the metal transport function of SLC39A8 in a different way.

SLC35F2-GFP localized to the plasma membrane and at the Golgi (**Fig. EV6E**). In cytometry assays, depletion of sorting nexin 2 (SNX2) in SLC35F2-GFP cells showed the strongest GFP reduction of all 134 protein-protein interactions tested (**Fig. 4D**). At the subcellular level, SNX2 depletion similarly reduced overall SLC35F2-GFP intensity (**Fig. EV6E**, quantified in **Fig. 6D**). SLC35F2 is the main importer of the experimental antitumor compound YM-155 (sepantronium bromide; (Winter *et al*, 2014)). In a viability assay based on transport of the toxic compound YM-155 by SLC35F2, we observed that expression of SLC35F2-GFP enhanced toxicity under drug treatment conditions, whereas depletion of SNX2 partially alleviated some of the effect, likely by decreasing the SLC35F2 levels at the plasma membrane (**Fig. EV6F**).

For 12/34 interactions of 23 SLCs localized to multiple compartments, SLC-GFP levels changed specifically at certain subcellular compartments (**Fig. 6D**). This included the organic anion transporter SLC22A11 (OAT4) and its interaction with the endoprotease FURIN (**Fig. EV6G**). Depletion of FURIN increased SLC22A11-GFP intensity specifically at the plasma membrane and not at the Golgi. Another example was the vitamin C transporter SLC23A1 localized in its GFP-tagged form at the plasma membrane and the lysosome (**Fig. EV6H**). While depletion of GalNAc transferase GALNT2 increased GFP intensity at the lysosome and at non-lysosomal regions (positive and negative for RFP, resp.), depletion of ubiquitin ligase SYVN1 predominantly increased lysosomal SLC23A1-GFP intensity (**Fig. 6D**), suggesting that ER quality control by SYVN1 is required for trafficking to the plasma membrane. Similarly, the copper importer SLC31A1 (CTR1) was found to be differentially regulated by two of its interactors. SLC31A1-GFP localised at the plasma membrane and in cytoplasmic vesicles. Depletion of syntaxin-7 (STX7) significantly increased both of these pools, while depletion of SLC31A1 interactor WDR6 increased its levels at the plasma membrane (**Fig. EV6I**, quantified in **Fig. 6D**).

Besides sorting nexins, proteins of the MAGUK (membrane-associated guanylate kinase) family were also strongly featured in the SLC-interactome, most notably associating with the amino acid transporter SLC43A2 (LAT4). In its conditionally expressed GFP-tagged form, SLC43A2 localized to the plasma membrane and to cytoplasmic vesicles (**Table EV3**). In our co-purification analysis (**Appendix Fig. S7**), prominent interactions emerged with the members of the LIN- and CASK/DLG1-complex (LIN2/CASK, LIN7/LIN7C, SAP97/DLG1 complex, CORUM (Tsitsiridis *et al*, 2023), and with MPP1 complex members (GPC, 4.1R/EPB41, p55/MPP1 complex, CORUM) (**Fig. 6F**, **Dataset EV2**). Proteins in both complexes are characterized by PDZ, SH3 or L27 interaction domains, are kinases or kinase-like proteins, and can interact with the actin or microtubule cytoskeleton to concentrate, polarize or recycle proteins at the plasma membrane (Alewine *et al*, 2007). These functions are thought to have emerged early in evolution to allow tissue formation (Baines et al. BBA 2014) and are disease-related in the context of brain development (CASK) and haematology (EPB41) (Dubbs *et al*, 2022). Interestingly, the interactions between the cytoplasmic C-terminal tail of SLC43A2 and the PDZ domains of LIN7C and MPP1 were modelled with medium confidence using AlphaFold (**Fig. 6H,I, Appendix Fig. S12**). When we depleted the protein complex subunits by RNAi, SLC43A2-GFP was affected differently depending on which PDZ-domain protein was targeted. While depletion of LIN7C and CASK increased plasma membrane associated SLC43A2 by almost 30%, depletion of MPP1 did not affect plasma membrane associated SLC43A2 (**Fig. 6D**). Instead, we found locally increased signals in cytoplasmic structures reminiscent of accumulating SLC43A2 in the Golgi (**Fig. 6G**). This suggested that we observed two distinct interaction hubs involved in trafficking: one containing the membrane-associated guanylate kinase MPP1, possibly involved in shuttling SLC43A2 from the Golgi to the plasma membrane, and another containing LIN7C and CASK, potentially increasing turnover of SLC43A2 at the plasma membrane (**Fig. 6J**).

In summary, we showed that many interactors identified in this study contributed to the function of SLC transporters, by influencing localization, protein levels or activity. Our findings can serve as a starting point for more in-depth research and for possible therapeutic modalities.

## Discussion

The ensemble of proteins encoded by the large number of human genes (1,500-2,000) estimated to govern the transport of chemical substances across all kinds of cellular membranes can be termed the human transportome (Huang *et al*, 2004). It can be reasonably assumed that most chemical integration of cells, organs and organisms with the respective environments is orchestrated by this transportome. Yet we lack answers to even some of the simplest questions about this interface: what do the individual gene products transport? How is their activity regulated? Does integration of function occur within each system? Characterization of the protein environment of any given protein should, without much prior knowledge on function, represent a valuable way to provide links to the biochemical and biological processes to which the gene product is involved. The protein partners shed light on the subcellular environment, participation to specific molecular machines, potential obligate and facultative partners and possible modifiers of activity, stability and localization (Marsh & Teichmann, 2015; Gavin & Superti-Furga, 2003).

Among the transportome, solute carriers represent the largest superfamily with roughly 450 members. As part of a larger effort to start elucidating the transportome systematically, we present here the first comprehensive protein interaction study of the human solute carrier superfamily. We generated tagged-versions of each SLC under a doxycycline inducible promoter, using codon-optimized sequences, and used an experimental AP-MS workflow tailored to TM-proteins. Throughout the study, we address commonly experienced limitations when studying TM-proteins, including expression of the tagged protein, lysis conditions, purification and elution of enriched complexes. To predict protein interactions in a reproducible manner, we devised a machine learning-based method which integrates common PPI-scores (including SAINT, CompPASS) with several quantitative and uniqueness features, thereby overcoming limitations of previous studies which relied solely on GFP-expressing controls or only a few other transporters (Rebsamen *et al*, 2015; Heinz *et al*, 2020). Together, these methodological developments allowed us to identify thousands of novel PPIs and generate the first comprehensive protein interaction landscape of SLCs.

Was the effort successful? We would argue that the effort has been technically successful, even if very labour-intensive, and that it represents a very significant resource for the community. Has the guilt-by-association concept allowed us to “deorphanize” many transporters? The simple answer is no. Unlike GPCRs or nuclear hormone receptors, SLCs are thought to be much more promiscuous, as the sheer number of diverse chemical substances entering the human body vastly exceeds the number of transportome genes. Moreover, the functional proteomics approach presented here is not suitable for assigning relationships to potential cargos directly. However, the combination of similarity clustering, enrichment analysis for SLC properties and subsequent analysis of cluster-specific PPI-networks allowed us to derive novel insights into functions of several SLC families. The similarity clustering used in our study is limited due to the heterogenous expression and localization of SLCs and the lack of reciprocal AP-MS experiments, inherently resulting in a low-density network. It was previously shown that high density networks are needed to de-convolute binary PPI-networks to protein complexes (Uliana *et al*, 2023; Buljan *et al*, 2020; Salokas *et al*, 2022). Similarity-based clustering on the interactome profile of each tagged protein is, on the other hand a rarely used approach for interaction proteomics, as it is strongly penalized by data sparseness. Despite these shortcomings, the clustering strategy of our AP-MS strategy was successful in positioning each SLC within a particular protein environment that provides important insights into its involvement in cellular and biochemical processes.

By detecting interactors across the SLC lifespan, the SLC-interactome can serve as roadmap to modulate targeting, trafficking, activity and abundance levels of SLCs through their interaction partners. This is of critical interest for diseases associated with pathogenic SLC transporter expression linked to folding-deficient variants (Wiktor *et al*, 2021; Ohtsubo *et al*, 2011), trafficking/targeting errors (Rogala-Koziarska *et al*, 2019) or proteostatic pathways (Colaco *et al*, 2023; Xu *et al*, 2016). We thus envision that the identified interaction partners may serve as drug discovery route to modulate SLC-activity indirectly, by rescuing folding and/or assemblies by correctors, pharmaco-chaperones and potentiators (Gautherot *et al*, 2012; Bhat *et al*, 2021, 2019).

Our validation efforts assess the value and quality of the SLC-interactome data set by testing subcellular localization and relative protein levels as a proxy for protein stability. We tested the functional impact of newly discovered interactions for 137 SLC-protein interactions, a large number for AP-MS studies.

Follow-up studies using endogenously modified proteins, such as shown in this study for MTCH2 and its interactors, in a cellular system expressing both interaction partners will be required to confirm the function of these interactions. Currently, scarcity of well-working antibodies against SLCs is a limiting factor. We have recently promoted the development of recombinant high-affinity binders directed against SLCs as alternative (Gelová *et al*, 2024). Using available antibodies, we were able to confirm that SLC6A15 and its interaction with KLHL36 occur at physiological SLC expression level. The sodium-dependent neutral amino acid transporter SLC6A15 and E3 ligase adapter KLHL36 could also be linked via disease susceptibility. While loss of SLC6A15 function was reported to be associated with depression (Kohli *et al*, 2011), increased KLHL36 expression has been tentatively correlated with suicide attempts (Han *et al*, 2023). KLHL36 SNPs have been linked to increased resilience in US army soldiers (Stein *et al*, 2019). Similar individual, or systematic investigations of such connections can become valuable tools for research and discovery.

Ectopic expression of bait proteins for AP-MS can also have unexpected benefits. βTrCP proteins play essential cellular roles such as cell cycle control, DNA damage repair, metabolism and signalling, and are considered both oncogenes and tumour suppressor genes. While their substrates have been studied in great detail using AP-MS (Kim *et al*, 2015; Dorrello *et al*, 2006; Low *et al*, 2014) and BioID (Coyaud *et al*, 2015), SLC16A6 has not previously been reported as an SCF^βTrCP^ substrate. Nevertheless, our data clearly suggest a physiologically relevant relationship between SLC16A6 and SCF^βTrCP^. Interestingly, SLC16A6 expression is confined to certain neuronal and reproductive human tissues and its expression is hardly detectable in common human cell lines, including most cell lines used in the RESOLUTE project (Karlsson *et al*, 2021). This suggests that screening approaches using ectopic expression may yield biologically relevant insights on protein homeostasis that are difficult to detect using only endogenous components.

Overall, we conclude that SLCs are extensively regulated by the cellular proteostatic machinery, affecting their levels and localisation.

There are some limitations to our study that need to be considered. First, we relied for extraction of SLC-complexes on NP-40/Ipegal as detergent, except for mitochondrial localized transporters, when we used digitonin. Previous interaction proteomics studies focusing on transmembrane proteins used DDM (n-dodecyl-β-maltoside) or C12E8 (octaethylene glycol monododecyl ether) to extract under native lysis conditions TM-localized complexes (Celis-Gutierrez *et al*, 2019). We used NP-40 to have comparable lysis conditions with other large-scale studies (Huttlin *et al*, 2021, 2015). Certainly, different detergents will result in different interactomes. Second, we focused our analysis on the consensus protein sequence for SLCs and interactors, therefore excluding isoforms. It was previously shown that SLC isoforms vary not only in their expression patterns regarding cell type or tissue, but they also can have different lengths of C- and N-terminus, which may affect SLC localization and interactions (Shirakabe *et al*, 2006; Mazurek *et al*, 2010; Yoo *et al*, 2020). Finally, because of the elaborate and conservative scoring of interaction partners, we might have misassigned true positive interaction partners wrongly as background. Benchmarking of our data set was hampered due to low coverage of SLCs in protein complex databases and considerable variability of SLC protein interactions reported in public PPI-databases.

A valuable additional layer in the characterisation of the molecular environment of transporters would be the assessment of lipid-protein interactions (Corradi *et al*, 2019). There are emerging data supporting the importance of phospholipid composition affecting activity of transmembrane transporters (Hresko *et al*, 2016). Recent developments in proteome-wide mapping of protein-metabolite interactions by LiP-MS and other method have great potential in this sense (Piazza *et al*, 2018). The public release of our data will further facilitate integration of SLC-protein interactions, likely to offer new appreciation of the modular organization of the interactome. Lastly, we think that it would be intriguing to study the SLC-interactome across different cellular states, thus using AP-MS to identify dynamic rewiring of the PPI-network upon changes of the transporter activity or in metabolic state.

Attractive possibilities include changes in media compositions, for example alternative carbon sources or concentrations of amino acids, and performing similar mapping campaigns across these different settings. Such experiments may lead to the identification of links between cellular signalling and modulation of transporter activity.

A searchable version of the SLC-interactome data set is available at https://re-solute.eu/resources/dashboards/proteomics/network/. The resource allows exploration of interactions, further filtering of interactions and integration of interactions obtained from BioGRID and CORUM to allow visualization of protein complexes.

Our study should contribute to the general understanding of the interactome of membrane-embedded proteins in human cells. Seminal previous studies have focused on a clinically relevant section of GPCR using yeast two-hybrid (Sokolina *et al*, 2017) or have been dedicated to individual baits or specific relationships. Thus, it can be expected that this work may represent a blueprint for the systematic mass spectrometry-based characterisation of the interactome of several other groups of multipass TM-proteins, such as transporters of the ABC, P-type ATPase and aquaporin groups, ion channels, mechanosensitive receptors and GPCRs (von Heijne, 2007). Because of the diversity of the biochemical and biophysical properties of the membranous environment, including the dimensional limitations to diffusion, it would not be surprising if protein-protein interaction principles established mainly with soluble proteins would be different for multipass membrane proteins. We are now using artificial intelligence-driven approaches to extract what expect to be interaction modules or “archetypes” that may allow to better predict interactions of membrane proteins generally.

## Materials and Methods

### Cell line generation

HEK 293 Jump-In T-REx cells (Thermo Fisher; RRID:CVCL_YL74) were co-transfected with pJTI R4 DEST CMV-TO pA vectors containing strepII-HA tagged (C-terminal) or HA-strepII tagged (N-terminal), codon-optimized SLC cDNA sequences and the pJTI R4 Int vector that encodes for the R4 Integrase according to manufacturer’s instructions (for all SLC cDNA sequences see **Table EV1**). For more information see accompanying manuscript (Wiedmer and Teoh et al, accompanying manuscript).

For validation assays, HEK 293 Jump In T-REx cells were co-transfected with the pJTI R4 Int vector and with pJTI R4 DEST CMV-TO pA vectors containing GFP-tagged SLC sequences, and a reference RFP separated by an internal ribosome entry (IRES) sequence. The RFP (mCherry or mRuby3) was preceded either by a myristylation tag for plasma membrane localization, four COX8 presequences for mitochondrial localization, the TMEM192 coding sequence for lysosome localization, or a myristylation tag with an in-frame upstream start codon for pan-cellular localization (GFP cell lines see **Table EV3**, localization tag determined based on (Meixner *et al*, 2020) and Goldmann et al, accompanying manuscript.

To generate point mutations in the SLC16A6 coding sequence at the phospho-degron motif DSGVELT (239–245), mutagenesis primers were designed using online tools (NEBaseChanger, NEB) and introduced into a pDONR plasmid using a Q5 site-directed mutagenesis kit (NEB). The resulting plasmids containing S240A, S240D, T245E, and T245N modified SLC16A6 were recombined in a Gateway LR reaction to express doxycycline-inducible SLC16A6-strepII-HA for validation experiments. To generate cells expressing FLAG-KLHL36, HEK 293T cells were transduced with a lentiviral vector expressing KLHL36 under an EF-1α promoter. The HAP1 HA-dTAG-MTCH2 knock-in single cell clone cell line was generated by microhomology-mediated end joining with the PITCh system as previously described (Bensimon *et al*, 2020).

### Cell culture and mitochondrial fractionation

HEK 293 WT OE cells were grown on 15 cm cell culture dishes in high-glucose DMEM medium supplemented with 10% FBS and 1% PenStrep at 37°C. SLC expression was induced by the addition of 1 μg/mL Doxycycline for 24 h. Depending on the expression levels 40, 80 or 160 million cells corresponding to two, four, or eight 15 cm plates per replicate were harvested.

For the standard AP-MS procedure, cells were scrapped in PBS, pelleted at 600 x g at 4°C for 10 minutes and frozen at -80°C until further usage. For mitochondrial localized SLC baits, a mitochondrial enrichment was performed. Scraped cells were transferred to a 50 mL falcon tube to pellet cells. Next, pelleted cells were re-suspended in freshly prepared isolation buffer (IBc, 10 mM Tris-MOPS, 1 mM EDTA-Tris, 200 mM sucrose, H_2_O, pH 7.4, supplemented with 1x PI-cocktail) and incubated for 10 minutes on ice. Lysis was achieved by sonication with a Branson digital sonicator (cycle time of 30 sec, 10% amplitude and 0.5 s on/off). The cell homogenate was centrifuged at 600 x g at 4°C for 10 minutes to remove un-lysed cells. The supernatant was collected and transferred to a 15 mL falcon tube and centrifuged at 10,000 x g at 4°C for 15 minutes. The supernatant was discarded, and the mitochondrial fraction was washed with 500 μL IBc followed by another centrifugation step. The mitochondrial fraction was frozen at -80°C until further usage.

### Affinity purification of SLC baits

We first affinity purified the C-terminal fusion protein, and for SLCs which failed (e.g. expression levels), we tested the N-terminal construct. We selected HEK 293 as the cellular model, as among 1,206 cell lines reported in the human protein atlas HEK 293 cells express an average number of SLCs (see **Appendix Fig. S13**). For SLCs localized at the plasma, lysosomal, Golgi Apparatus (Golgi), Endoplasmic Reticulum (ER) or in vesicular membranes, cell pellets were lysed in freshly prepared lysis buffer containing 50 mM HEPES pH 8.0, 150 mM NaCl, 5 mM EDTA, 0.5% NP-40, 1 mM PMSF, 1x PI- cocktail, avidin (1 μg/mL), 1x phosphatase inhibitor cocktail 2 and 1x phosphatase inhibitor cocktail 3. The ratio of cells to lysis buffer was kept constant (2 plates = 1.8 mL lysis buffer). Lysates were clarified (14,000 x g at 4°C for 20 min) before they were incubated with 150 μL pre-equilibrated StrepTactin Sepharose (50% suspension) beads for 2 h at 4°C. Beads were gently centrifuged at 1,000 x rpm for 1 min at 4°C and the flowthrough was discarded. Beads were then washed twice with 1 mL lysis buffer and centrifuged. Next, beads were resuspended in 1 mL of washing buffer (50 mM HEPES pH 8.0, 150 mM NaCl, 5 mM EDTA) and transferred to BioSpin membrane columns (Bio-Rad). Beads were washed 3x with 1 mL washing buffer. Finally, proteins were eluted by incubation of beads with 125 μL 2% v/v SDS in washing buffer for 15 min.

For mitochondria-localized SLCs the enriched mitochondrial pellets were resuspended in 3 mL freshly prepared lysis buffer with the same composition as above except that instead of NP-40, 1% digitonin was used as detergent. To increase protein extraction yields, samples were briefly sonicated in a Branson sonicator with microtip (setting: 10 s sonication time, 0.5 s on/off cycles with 10% amplitude). After this step, mitochondrial samples were handled as outlined above for SLCs localized at other cell membranes.

Detailed descriptions of the AP-MS protocols were made publicly available under https://doi.org/10.5281/zenodo.7457416 and under https://doi.org/10.5281/zenodo.7462207 for the mitochondria-localized SLCs.

### Sample preparation for AP-MS

Sample preparation was performed by a single-pot solid-phase-enhanced sample preparation (SP3) procedure following an adapted protocol (Müller *et al*, 2020). In short, eluted proteins were reduced with DTT (10 mM) and incubated on a shaker at 56°C at 600 rpm agitation for 1 h. Cysteine residues were alkylated with IAA (50 mM) followed by incubation in the dark at RT for 30 min. 8.8 μL of Mag- beads prepared according to manufacturer’s instructions were added. Next, proteins were precipitated with 300 μL Acetonitrile (ACN) and the resulting solution was incubated for 10 minutes on a shaker followed by a 10-minute incubation on the bench. Beads were magnetized and the buffer was removed, after which three washing steps with 200 µL 80% (v/v) ethanol with 2 min incubation in-between followed. Next, beads were washed twice with 180 μL ACN with 2 min incubation between washings steps. For digestion, beads were resuspended in 100 μL 50 mM Tris-HCl (pH 8) with 1 μg Trypsin/Lys-C Mix (37°C at 600 x rpm on). Samples were acidified with 2 μL of 30% TFA in H_2_O and after magnetising beads the supernatant was collected and centrifuged for 1 min at 14,000 x g to remove remaining beads. Peptides were purified by solid phase extraction C18 stage-tips protocol. After washing, peptides were eluted with 2x 50 μL of 60% ACN, 0.1% TFA. Samples were dried under reduced vacuum at 45°C and stored at -20°C until analysis. Peptides were reconstituted in 30 μL MS-buffer of 0.1% TFA. A more detailed procedure for the MS-acquisition can be found under: https://zenodo.org/records/7462253.

### AP-MS data acquisition

Mass spectrometry data were acquired on a Q Exactive Orbitrap MS coupled to a Dionex Ultimate 3000 RSLCnano system interfaced with a Nanospray Flex Ion Source. Peptides were loaded onto a trap column (Pepmap 100 5 μm, 5 x 0.3 mm) at a flow rate of 10 μL/min 0.1% TFA. After loading, the trap column was switched in-line with the analytical column (50 cm, 75 μm inner diameter analytical column packed in-house with ReproSil-Pur 120 C18-AQ, 3 μm) kept at 40°C to which an ESI Emitter Fused Silica was fitted. Data acquisition was conducted at a flow rate of 230 nL/min with a 120 min gradient (4% to 24% solvent B (90% ACN 0.4% FA) in 86 min, 24% to 36% solvent B within 8 min and 36% to 100% solvent B within 1 min, 100% solvent B for 6 min, 100% to 4% solvent B in 1 min and 4% solvent B for 18 min). Eluted peptides were ionized in a positive mode (1.8 kV). MS analysis was performed in a data-dependent acquisition (DDA) mode. Full MS scans were acquired in a precursor mass to charge (m/z) range of 375–1650 m/z in the orbitrap at a resolution of 70,000 (at 200 Da). Automatic gain control (AGC) was set to a target of 1x 10^6^ with a maximum injection time of 55 ms. Precursor ions were selected by a Top10 approach using a quadrupole isolation window width of 1.6 Da and higher energy collision induced dissociation (HCD) at a normalized collision energy (NCE) of 28%. The MS2 AGC target was set to 1x 10^5^ with a maximum injection time of 110 ms and an orbitrap resolution of 17,500 (at 200 Da). Dynamic exclusion for selected ions was set to 40 s. A single lock mass at m/z 445.120024 and XCalibur version 4.1.31.9 and Tune 2.9.2926 were used.

Part of the AP-MS data were acquired on a Q Exactive HF-X Hybrid Quadrupole-Orbitrap MS. Peptides were separated by reverse-phase chromatography using a nano-flow HPLC (Ultimate 3000 RSLC nano system). Per injection, 3 μL sample was loaded onto a trap column (PepMap C18, 5 mm x 300 μm ID, 5 μm particles, 100 Å pore size) at a flow rate of 10 μL/min using 0.1% TFA. The trap column was switched in-line with the analytical column (PepMap C18, 500 mm x 75 μm ID, 2 μm, 100 Å) for analysis. During the next 10 min a flow rate of 50 μL/min with 0.1%FA 70% Methanol was applied. Elution was achieved at constant T of 40°C (external butterfly heater controlled by column heater controller). For separation, solvent A with 0.4% FA and organic solvent B with 0.4% FA 90% ACN were used. Flow rate was set to 230 nL/min and a 120 min gradient was used (4% to 24% solvent B within 81 min, 24% to 36% solvent B within 8 min and 36% to 100% solvent B within 1 min, 100% solvent B for 6 min before equilibrating at 4% solvent B for 18 min). Eluted peptides were ionized in a positive mode (1.8 KV) using an Easyspray nanospray Source.

Data acquisition was performed in a DDA mode. Full MS scans were acquired with a m/z range of 375– 1650 m/z in the orbitrap at a resolution of 60,000 (at 200 Da). AGC was set to a target of 1 x 10^6^ with a maximum injection time of 55 ms. Precursor ions for MS2 analysis were selected using a Top10 approach using a quadrupole isolation window of 1.6 Da and HCD at a NCE of 28%. AGC target was set to 1 x 10^5^ with a maximum injection time of 55 ms and an orbitrap resolution of 30,000 (at 200 Da). Dynamic exclusion for selected ions was 15 s. A single lock mass at m/z 445.120023 was employed for internal recalibration during the run. XCalibur version 4.3.73.11 and Tune 2.13 Build 3162 were used to operate the instrument.

### Processing of AP-MS

MS raw files were converted to mzML files with MSConvert (3.0.21128-7376ae988) (Chambers *et al*, 2012). Acquired spectra were searched using Philosopher’s pipeline mode with MSFragger (da Veiga Leprevost *et al*, 2020). The identification search was performed against the canonical human proteome obtained from UniProtKB (downloaded on the 2021-04-01, status reviewed). The database was supplemented with 48 lab contaminants. The search parameters were set to include fully tryptic peptides, allowing for up to two missed cleavage-sites, with carbamidomethylation on cysteine residues as static modification. Further up to five variable modifications including oxidation on methionine and N-terminal acetylation were enabled. Mass tolerance was set to 50 ppm for precursor ions and 20 ppm for fragment ions. The peptide length was restricted from five to 63. A MS1 m/z- range from 375 to 5000 m/z and precursor charges states +1 up to +4 were set. Peptide assignment validation was performed with PeptideProphet and protein inference with ProteinProphet. FDRs were controlled at 1% peptide FDR and 1% protein FDR respectively. Files were searched one at a time similar to other large-scale interaction proteomics studies (Huttlin *et al*, 2021).

The assembled bait-prey matrix for each data modality served as input for feature generation (**Fig. 1D**). The 1,432 MS-injections covering 358 SLCs and 28 GFP controls prepared with the standard protocol and measured on the QE were assembled. For the QE HF-X matrix, the data of 23 SLCs and 4 GFPs (107 MS-injections, 1 technical injection was missed) were combined. Lastly, the mitochondrial localized SLCs bait-prey matrix was assembled, containing 208 MS-injections of 47 SLCs and mitochondria targeted GFPs. The data were independently assembled to account for the difference in the background. Next, the data per bait were grouped together, and reproducibility filtering strategy was employed, to remove sparse or wrongly assigned protein identifications, thereby lower protein FDR below 1%. The mean SPC per biological replicate was derived only for proteins which were quantified across technical injections. After this filtering step, the protein FDR was re-estimated by counting passing decoys, resulting in 0.64% FDR for the standard QE modality (6,962 proteins, 45 decoys), 0.14% FDR for the mitochondrial modality (3709 proteins, 5 decoys), and a 0.82% FDR for the QE HF-X modality (5,434 and 45 decoys).

### Assembly of feature matrix for scoring of PPIs

First, a set of scoring features was generated. Interaction scores for each protein were generated using CompPASS (Sowa *et al*, 2009) and SAINTexpress (Teo *et al*, 2014). As inputs, the bait-prey matrix per data modality was used. For CompPASS the normalization factor was set to 0.95. For the SAINTexpress scoring, we generated 100 *in silico* controls, specific for each SLC. To achieve this, we randomly sampled SPCs for each unique protein identifier from SLCs and GFP controls, only considering SLCs with similar summed up signal across all proteins. For this we ranked the absolute delta difference of the mean SPC to other SLCs and GFPs within each data modality. For the random sampling, only data from AP-MS of SLCs with similar summed up signal were considered, resulting in randomized controls. For the standard QE modality, 20% of all samples were used for random sampling, whereas for the mito and QE-HFX data sets, which had fewer samples, up to 40% of all samples were used. For the sampling the abundance of the SLC bait protein was excluded. These controls were used for SAINTexpress scoring with default parameters.

Second, a set of quantitative, annotation and experiment wide features was derived (see **Table EV2** for a list of all features). For each protein pair of each SLC, additional quantitative features were derived, including the normalized spectral abundance factor (NSAF) (Neilson *et al*, 2013) and the ratio of proteotypic to total spectral counts. Further, data set wide features per interaction partner were added, including the sum of total SPC and the ratio of total SPC against the data-wide summed up signal. Additionally, SLC-protein pair feature was derived, including the FC against GFP (per data modality), and an additional FC against the SPC of each protein of all other SLC and GFP samples (leaving out the signal in the SLC experiment itself for which the FC was derived). As sample specific quantitative features, the summed mean spectral counts, the number of quantified proteins, and the SLC bait abundance of each sample were added. Finally, as deterministic features, the experimental protocol (‘standard’ or mitochondrial AP-MS) and the MS-platforms were added. Finally, all data modalities were assembled (**Dataset EV4**).

### Curation of labels for SLC protein interactions

For the ML-based probability scoring of interactions, two sets of labelled PPIs were generated. The first set contained overlapping PPIs with BioGRID (v.4.4.223) (Oughtred *et al*, 2021), filtered to contain only the experimental category identifiers “Affinity Capture-MS”, “Affinity Capture-Western”, “Reconstituted Complex” and “Co-crystal Structure”. All matched pairs were set as true. For the second set, we used SLC-protein pairs reported in PDB. This list was expanded by including bait self-loops and curated interactions. The curated PPIs were selected considering PPIs recovered within the SLC-interactome and BioGRID, taking the abundance and specificity of each SLC-protein pair into account. The final feature matrix contained 693 true and 970 false labels.

### ML-based scoring of PPIs

The aim of the ML learning was to integrate the features (see above) in one model to calculate an interaction probability for each SLC-protein combination. First the features were pre-processed. To map all input features to real numbers, each feature was either log- or z-transformed, or taken as-is. Non-finite values were imputed to complete the feature matrix. To assure numerical stability during model fitting, the transformed features were subsequently normalized to have zero mean and unit standard deviation. To provide reliable probabilities whether a protein is an interactor, an ensemble of 30 Radial Basis Function (RBF) based classifiers is fitted iteratively on different random subsets of all speculatively labelled interaction partners. Using speculative labels allows for potential mistakes in database entries and is considered in model fitting by allowing for BioGRID based reports of interaction between interactors and SLC as well as lack of reported interaction to be wrong. We assume however that a small set of exquisitely curated interaction states can be trusted without doubt. While always trusting the curated labels, model fitting redraws novel BioGRID-derived interaction labels according to the RBF based interaction probabilities. This process is repeated until the respective labels show less than 1% differences in two consecutive runs. Against the curated PPI data set averaged ensemble predictions reached an accuracy of 95%, with a sensitivity of 0.89 and specificity of 1.0. We used a combination of thresholds (RBF probability ≥ 0.4, log2FC against GFP > 2, FC against other samples > 1.5, entropy filter < 0.75, quantification frequency < 99%) to classify proteins as interactors or background. From the initial 634,125 protein pairs roughly 2.99% or 18,991 interactions were assigned as PPIs.

### Analysis of SLC16A6 phospho-mutant AP-MS data set

AP-MS data of SLC16A6 phospho-mutants were generated as described. Data were searched together with the original SLC16A6^WT^ AP-MS raw files and 24 GFP controls using Philosopher. Peptides were quantified with freequant (default parameters). For further analysis, the peptide level output was used. The SLC16A6 phospho-mutants and SLC16A6^WT^ samples were filtered and the Top3 most intense peptides across the experiment were used to infer protein abundances. Protein abundances were normalized using total signal scaled to the median signal across the experiment. Next, abundances were log2 transformed, and missing values were imputed sampling from a normal distribution around the lowest 5% quantile of all abundances. For each phospho-mutant, the log2FC against the SLC16A6^WT^ was derived. A log2FC of ±1 was used to assign enriched or depleted interactors respectively. The results are reported in **Dataset EV3**.

### Curation of a PPI-reference library of SLCs

To benchmark our SLC-interactome, we assembled a reference PPI-library covering multiple PPI- databases. For this we mined BioGRID (Oughtred *et al*, 2021), IID (downloaded on the 11.05.2021) (Kotlyar *et al*, 2022), IntAct (Orchard *et al*, 2014) (downloaded on the 11.09.2023) and STRING v12 (Szklarczyk *et al*, 2023). BioGRID was filtered to include PPIs from the categories "Affinity Capture-MS", "Affinity Capture-Western", "Reconstituted Complex" and "Co-crystal Structure”. IID was limited to experimental data and human proteins. PPIs without an associated reference were removed from IID and experimental methods were limited to AP-MS. STRING was filtered to cover physical interactions with a confidence threshold of 0.4. IntAct was filtered to contain only human proteins in combination with the method identifiers: “MI:0006”, “MI:0007”, “MI:0019”, “MI:0096”, “MI:0114” and “MI:0676”. For the reference count, interactions obtained from STRING were not considered. For categorization of proteins containing TM-domains the UniProtKB database annotations for topological domain “transmembrane” (downloaded on the 2022-05-13) was used.

### Benchmarking of SLC-interactome

We benchmarked our data set on PPI and protein complex levels. On PPI-level, we performed the benchmark against BioPlex (Huttlin *et al*, 2015, 2017, 2021) and HuRI (Luck *et al*, 2020), whereas on complex level we used the CORUM complex (Tsitsiridis *et al*, 2023) database, filtered for human complexes and heteromers. CORUM complexes were translated to a PPI-network assuming full connectivity of all subunits. Next, we supplemented interactions between all interactors for each SLC and generated a supplemented PPI-network. This network was used to calculate the overlap with CORUM interaction pairs. To estimate if CORUM interaction pairs were enriched within the original SLC co-purification PPI-network, we permutated network nodes. For permutation, we sampled from all potential interactors and SLCs present in our data set but conserved the network topology. This permutation was repeated 10,000 times, and the p-value was derived by a rank-based test. Next, we used the CORUM data to derive counts of complexes and subunits present in the SLC-interactome. For counting complex-subunits, we considered subunits present in multiple assemblies. CRAPome 2.0 database (Mellacheruvu *et al*, 2013) was used to retrieve proteins regularly found in AP-MS negative controls.

### Construction of the SLCome

To construct a structure-based phylogenetic tree of the human SLCs, we retrieved the similarity matrix of protein structure models from (Ferrada & Superti-Furga, 2022), restricted to the 446 human SLCs which the RESOLUTE data set studied. SLC22A20P was not included in the structural analysis. Distances were obtained by subtracting similarities from 1 and used to cluster SLC structures hierarchically (hclust function in R, version 4.3.3, with Ward’s method). An unrooted tree was constructed using square root scaled branch lengths (ggraph library, version 2.2.1). Node positions were adjusted manually to reduce overlaps and optimize overall arrangement. In the final unrooted phylogenetic tree, which we termed SLCome, edges were colour-coded by structural folds (Ferrada & Superti-Furga, 2022) and SLC nodes were coloured if they were part of the SLC interactome. The number of novel and reported interactions were reported for each structural fold, using the reference PPI-library assembled for this study.

### Prediction of structures of protein interactions by AlphaFold

We used AlphaFold2 multimer (version 2.3) (Evans *et al*, 2021) to model binary PPIs. For each prediction at least 25 models were generated. Amber was set to TRUE. Predicted structures were assessed with the pDockQ-score similar to another study (Burke *et al*, 2023). A pDockQ score > 0.5 was considered as high-confidence model, whereas a pDockQ < 0.5 but > 0.23 were classified as medium confidence structure. A pDockQ score < 0.23 was considered as low confidence structure. Negative SLC-chaperone interaction pairs for structural modelling were randomly sampled considering the lowest 25% quantile of quantitative signal across our data. Statistics were obtained using an unpaired two-sided t-test. Structures were visualized with UCSF ChimeraX 1.7.1 (Pettersen *et al*, 2021), by selecting the structure with the highest pDockQ.

### Clustering analysis of the SLC-interactome

To cluster SLC-interactomes, we performed hierarchical clustering on the Jaccard distance of all interactors (hclust function in R, with Ward’s method). The selected number of clusters (k=38) was identified based on the mean silhouette width (**Appendix Fig. S5)**. GO enrichment analysis against GO Biological Process 2023 (GO:BP:2023) was performed with Enrichr (Kuleshov *et al*, 2016) for individual clusters. Of the 1,986 unique terms, only 25% were found in more than three clusters. This resulted in 681 terms used for hierarchical clustering (Euclidean distance, hclust function in R, with Ward’s method, k=44 **Appendix Fig. 6A**). For the identification of representative GO terms covering major biological functions, we used GO semantic similarity analysis with the R-package GoSemSim (Yu, 2020). Similarities were calculated per cluster using the relevance method. Subsequently, we used the reduce function to identify parental terms per cluster (score=size, cutoff of 0.7). The selected terms were visualized as functional cluster annotations in **Fig. 3A**. Similarities between enriched GO terms were derived by using the Wang method with a similarity cutoff of 0.65 (**Fig. 3B**).

To classify protein functions associated with the SLC interactome in **Fig. EV4A**, we used the gene set enrichment tool Enrichr and the gene set GO:BP:2023. The top ten terms were ranked by adjusted p-value and consolidated into three major categories. P-values were aggregated using P Fisher’s combined probability test.

To classify subcellular protein localizations associated with the SLC interactome in **Fig. EV6**, we used Enrichr and the gene set GO Cellular Component 2023 (GO:CC) with individual SLC interactomes. To consolidate terms into ten subcellular localizations (**Fig. EV6A)**, we generated curated lists of GO:CC terms for each subcellular localization and sequentially and conditionally evaluated each term or its parental terms for association with these subcellular localizations. P-values were consolidated as outlined above for GO:BP with a p-value cutoff < 0.05.

### SLC property enrichment analysis

As a measure of functionality, we tested clusters for enrichment of SLC functional properties of different classes (coupled ion, family, fold, location, substrate class). Respective classes and annotations are described in Goldmann et al, accompanying manuscript. Fisher’s exact test for overrepresentation was performed for each functional property in each cluster, only considering properties with at least 3 annotations. Resulting p-values were corrected for multiple testing using the Benjamini-Hochberg procedure for each class separately, and enrichments of functional SLC properties in clusters were called at 20% FDR.

### Protein stability and subcellular localization assay

For all protein stability and subcellular localization assays, cells were passaged onto poly-L-lysine coated and tissue culture treated 96 well plates at 10,000 and 25,000 cells per well in DMEM supplemented with 10% FBS, 1% penicillin/streptomycin and 1 µg/mL doxycycline 24 hours before analysis. For flow cytometry, cells were PBS washed, trypsinized, and resuspended in FACS buffer (5% FBS, 1 mM EDTA in PBS) before analysis on a BD LSRFortessa cytometer. Signals were acquired for GFP, RFP, and BFP as outlined in the Reagent and Tool Table. Cytometry analysis was performed using FlowJo by gating for healthy and single cells (usual range 5,000–10,000 per sample) and additionally for BFP positive cells in the case of cDNA overexpression. Median fluorescent values were processed using python software. Significant changes had to fulfill three criteria: (1) The mean of the median GFP:RFP ratio and the mean of the median GFP values had to change by at least 10% compared to the respective mean of the median parameter following control RNAi; (2) Statistical significance was defined as a p-value less than 0.01 in an independent t test comparing GFP:RFP ratios and of GFP values between control and interactor RNAi; (3) The change in GFP values, relative to the mean of the median GFP values after control RNAi, had to occur in the opposite direction to the change in RFP values, relative to the mean of the median RFP values; or the mean change in RFP values had to be no more than half of mean GFP value change.

For high-throughput imaging, media were exchanged with fresh DMEM containing 10% FBS, 1% penicillin/streptomycin and 1 µg/mL Hoechst 33342 and imaged on an Opera Phenix with 40x water objective and signal acquisition for Hoechst, GFP and RFP as outlined in the Reagent and Tool Table. Image analysis was performed using Fiji (Schindelin et al. Nature Methods 2012) with custom macros to measure signal Pearson correlation and thresholded signal areas, intensities and overlaps (BioImage Archive submission S-BIAD1105) and processed using python software. Significant changes had to fulfill three criteria: (1) For SLCs residing at a single subcellular location and for the cell body reference RFP, as well as for GFP signals at the reference RFP, the mean GFP intensity had to change by more than 10% compared to the mean GFP intensity after control RNAi, and more than 25% for GFP signals not overlapping with the reference RFP; (2) Statistical significance was defined as a p-value less than 0.01 in an independent t test comparing GFP intensity between control and interactor RNAi; (3) For SLCs residing at a single subcellular location and for the cell body reference RFP, as well as for GFP signals at the reference RFP, the mean GFP intensity change had to occur in the opposite direction to the mean RFP intensity change at the reference RFP. For interactions that significantly changed multi-location SLCs at a specific subcellular location, two additional criteria were applied: (1) Statistical significance was defined as a Benjamini/Hochberg corrected p-value less than 0.001 in an independent t test comparing relative GFP intensity changes between GFP positive pixels that overlapped with, and those that did not overlap with, the reference RFP; (2) the mean GFP intensity change at reference RFP pixels had to occur in the opposite direction to the mean GFP intensity change at non-reference RFP pixels, or the difference between the GFP intensity changes had to be at least one-third of the absolute value of the larger change.

For drug treatment, cells were plated on treated 96 well plates at 10,000 cells per well 2 days before analysis and induced with doxycycline-containing media 24 hours before analysis. An additional 10 μL drug-containing media (0.5 μM TAK-243; 0.5 μM MLN4924; 100 nM Bafilomycin A1; or 0.05% DMSO in full media) were supplemented 4.5-6 hours before analysis.

For RNAi-mediated depletion, cells were first seeded on tissue culture treated 6 well plates at 200,000 cells per well a day prior, or at 400,000 cells per well on the day of transfection in antibiotics-free media. A transfection mix of 400 μL OptiMEM with 1 μL DsiRNA (Integrated DNA Technologies; stock concentration 20 uM prediluted in duplex buffer from three individual DsiRNAs per gene) and 4 μL Lipofectamine RNAiMAX Transfection Reagent (Thermo Fisher Scientific) was incubated for 10 minutes before dropwise addition to cells and incubation for 48 hours. Cells were trypsinized, counted and seeded with doxycycline-containing media on 96 well plates with 8 wells per sample. To assess protein stability by cytometry, 4 wells were treated with DMSO or inhibitors to ubiquitination, neddylation or lysosome acidification 4.5–6 hours before analysis. See additional details regarding DsiRNA preparation, validation by RT-qPCR and DIA-based profiling of RNAi treated cells in **Appendix and Appendix Fig. S11, S12, Table EV5**, and **Datasets EV5 and EV6.**

Interactor cDNA was obtained from Addgene, from Novartis or ordered from Genscript (see **Table EV4** for details including references and sequences). Coding sequences were cloned into Gateway^TM^ pDONR^TM^221 (Thermo Fisher Scientific) and recombined into a constitutive expression vector containing an EF-1α promoter, the interactor cDNA, an IRES and a TagBFP-NLS sequence to control for expression. For cDNA overexpression, HEK 293 cells were first seeded on tissue culture treated 12 well plates at 100,000 cells per well a day prior, or at 200,000 cells per well on the day of the transfection. Untagged coding sequences were expressed from a constitutive EF1alpha promoter followed by a BFP-NLS reporter separated by an IRES. Transfection was performed by mixing 50 μL OptiMEM with 3 μL Lipofectamine 3000 (Thermo Fisher Scientific), 5 minute incubation, and mixing the Lipofectamine solution with 50 μL OptiMEM supplemented with 1 μg plasmid DNA and 2 μL Lipofectamine P3000 reagent before dropwise addition to cells. See Appendix **Table EV4** for sequences and references of interactor cDNAs. All assay results are summarized in **Table EV6.**

### Cadmium uptake assay

To assess changes in SLC39A8 transporter activity, a combination of a cadmium uptake-assay with RNAi of interactors was used. Experiments were conducted in HEK 293 WT OE cell lines applying the same conditions as described for the protein level and localization assays. After 48 h RNAi treatment, cells were seeded on 384-well plates and expression of SLC39A8 was induced for 24 h. Next, cells were washed six times with 20 μL uptake buffer using AquaMax DW4 (AquaMax DW4 Microplate Washer), leaving 20 μL residual buffer. 20 μL of component A (FLIPR® Calcium 5 Assay Kit) were added, and plates were centrifuged for 1 min at 1,000 x g followed by 2 h incubation at RT in the dark. The assay was performed on an FDSS 7000EX with an excitation wavelength of 480 nm and an emission of 540 nm with an exposure time of 200 ms for 20.5 min. After 50 sec, 20 μL uptake buffer (117 mM NaCl, 4.8 mM KCl, 1 mM MgCl_2_, 10 mM glucose, 10 mM HEPES pH 7.4) was added. Next, cells were incubated with 5 μM CdCl_2_ in uptake buffer at second 351 for 15 min and images were acquired every 2 seconds. Edge effects were mitigated by omitting border columns and rows. The mean fluorescence signal across replicates (Biological replicate 1 had 12 wells whereas biological replicate 2 and 3 each 8 wells) were derived and min-max scaled. The mean AUC was calculated by the trapezoid rule. Significance was tested across mean AUC using an unpaired t-test. Cell viability was measured by CellTiter-Glo (Promega). Results are summarised in **Appendix Fig. S9** and **Dataset EV7**.

### TRACT assay

HEK 293 Jump In T-REx cells expressing SLC1A3, SLC1A2 or SLC22A3 were assessed by Impedance-based Transport Activity through receptor Activation (TRACT) assays using MP real-time cell analyser (RTCA) as described previously (Sijben *et al*, 2022). Selected interaction partners were treated with RNAi as described above. SLC22A3-mediated uptake of MPP^+^ was measured using xCELLigence MP RTCA as described previously (Mocking *et al*, 2022). The methods and assay conditions are described in more detail in the Appendix Methods.

## Data availability

The MS data have been deposited to the ProteomeXchange Consortium via the PRIDE partner repository with the data set identifiers PXD055605 for the SLC-interactome, PXD051747 for proteome profiling of cell lines, PXD055192 proteome profiling of RNAi validations and PXD055141 for the SLC16A6 mutant data set. Predicted structures of SLC-protein interactions are available at <u>10.5281/zenodo.13693141</u>. Subcellular localization data containing images, results and macros are available at the BioImage Archive under submission S-BIAD1105. Protein stability data containing cytometry files are available at Zenodo <u>10.5281/zenodo.12758951</u>. Protein interactions and networks can be viewed on an interactive dashboard (https://re-solute.eu/resources/dashboards/proteomics/network/).

## Author contribution

**Fabian Frommelt:** Conceptualization; Data curation; Formal analysis; Investigation; Methodology; Project administration; Supervision; Validation; Visualization; Writing – original draft; Writing – review & editing. **Rene Ladurner:** Conceptualization; Data curation; Formal analysis; Investigation; Methodology; Supervision; Validation; Visualization; Writing – original draft; Writing – review & editing. **Ulrich Goldmann:** Conceptualization; Data curation; Formal analysis; Funding acquisition; Methodology; Visualization; Writing – review & editing. **Gernot Wolf:** Investigation; Methodology. **Alvaro Ingles-Prieto:** Investigation; Methodology. **Eva Lineiro-Retes:** Data curation; Investigation; Methodology. **Zuzana Gelová:** Investigation; Validation. **Ann-Katrin Hopp:** Visualization. **Eirini Christodoulaki:** Formal analysis; Visualization. **Shao Thing Teoh:** Visualization; Writing – review & editing. **Philipp Leippe:** Visualization; Writing – review & editing. **Manuele Rebsamen:** Investigation. **Sabrina Lindinger:** Data curation; Investigation; Validation. **Iciar Serrano:** Data curation; Investigation; Validation. **Svenja Onstein:** Investigation; Methodology. **Christoph Klimek:** Investigation; Methodology. **Barbara Barbosa:** Investigation; Methodology. **Anastasiia Pantielieieva:** Data curation. **Vojtech Dvorak:** Visualization. **J. Thomas Hannich:** Investigation; Supervision. **Julian Schoenbett:** Data curation; Validation. **Gilles Sansig:** Data curation; Validation. **Tamara A. M. Mocking:** Data curation; Formal analysis; Validation. **Jasper F. Ooms:** Validation. **Adriaan P. IJzerman:** Funding acquisition; Resources; Supervision. **Laura H. Heitman:** Funding acquisition; Resources; Supervision. **Peter Sykacek:** Formal analysis; Methodology; Resources; Writing – review & editing. **Juergen Reinhardt:** Conceptualization; Funding acquisition; Resources; Supervision. **André C Müller:** Conceptualization; Funding acquisition; Methodology; Resources; Supervision. **Tabea Wiedmer:** Conceptualization; Funding acquisition; Project administration; Supervision; Writing – original draft; Writing – review & editing. **Giulio Superti-Furga:** Conceptualization; Funding acquisition; Methodology; Project administration; Resources; Supervision; Writing – original draft; Writing – review & editing.

## Acknowledgments

This study received funding from the RESOLUTE consortium. RESOLUTE has received funding from the Innovative Medicines Initiative 2 Joint Undertaking under grant agreement No 777372. This Joint Undertaking receives support from the European Union’s Horizon 2020 research and innovation programme and EFPIA. The last year of work, including validation of data and writing of the manuscript was supported mainly by the Austrian Academy of Sciences. This article reflects only the authors’ views and neither IMI nor the European Union and EFPIA are responsible for any use that may be made of the information contained therein. G.S-F. was supported by the Austrian Academy of Sciences throughout. We thank the CeMM Proteomics-Metabolomics facility and the IMP (Research Institute of Molecular Pathology) proteomics facility for support in MS data acquisition, especially Andrea Rukavina for the full proteome profiling of RNAi treated samples and especially Elisabeth Roitinger and Gabriela Krssakova from the IMP for MS-data acquisition support. We thank Klaus Kratochwill from the Medical University of Vienna for providing access to the mass spectrometer throughout the project. We want to thank Georg Winter and lab members, especially Natalie Scholes, for assistance in setting up the protein stability assay pipeline. Stefan Kubick and the CeMM Molecular Discovery Platform (MDP) team members for support with the screening equipment and CeMM IT team, especially Patricia Carey for their support. We further want to thank Anthony Orth at Novartis Pharma AG BR DSc for providing interactor cDNA constructs. The prediction of structures by AlphaFold have been achieved using the Vienna Scientific Cluster (VSC). We want to thank Ariel Bensimon for providing the HAP1 cell line expressing MTCH2. We thank Gabriel Onea for critical reading and feedbacking the manuscript.

## Supplementary information

### Tables

- **Table EV1**: SLC baits used for generation the interaction data.
- **Table EV2:** Features used for ML-based prediction of interaction probability.
- **Table EV3:** GFP cell line summary used for validation of SLC-protein interactions.
- **Table EV4:** DsiRNAs used for validation of SLC protein interactions.
- **Table EV5:** PCR primers for validation of selected RNAi pools by RT-qPCR.
- **Table EV6:** Summary of validation results.
- **Table EV7**: DsiRNA Design Identifiers

### Datasets

- **Dataset EV1:** Protein interaction partners identified within the SLC-interactome.
- **Dataset EV2:** Protein stability and subcellular localization assay results.
- **Dataset EV3:** AP-MS of SLC16A6 mutants.
- **Dataset EV4:** Assembled features of all PPI-pairs.
- **Dataset EV5:** RNAi validation by qPCR.
- **Dataset EV6**: Full proteome validation profiling after RNAi treatment.

• **Dataset EV7**: Cadmium uptake assay result for SLC39A8.

---

**Figure EV1.**
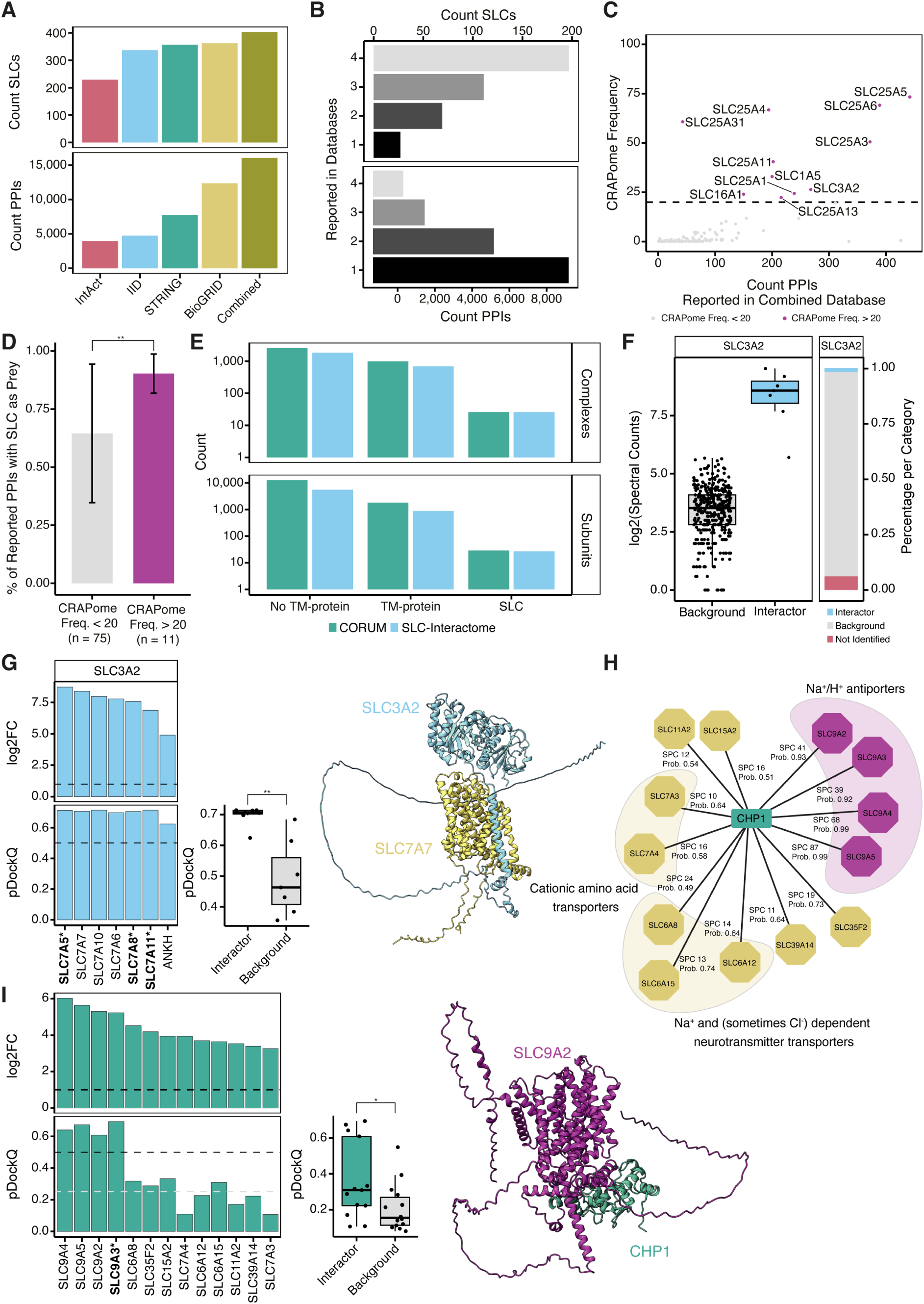
Characterization of SLC-protein interactions, SLCs as background and SLCs in complexes. **(A)** SLCs and PPIs reported across protein interaction databases. **(B)** Overlap of SLCs and PPIs across protein interaction databases. **(C)** Reported PPIs across the PPI-library were plotted against frequency of identification in the CRAPome database. 127 SLCs were reported in the PPI-library and were also part of CRAPome (CRAPome frequency > 20% in violet, < 20% in grey). **(D)** Comparison of the share of PPIs reported in the prey role for SLCs in BioGRID with a CRAPome frequency > 20% (n=11) against the share of PPIs reported in the prey role for SLCs with a CRAPome frequency < 20% (n=75). The two groups showed a significant difference (Two sample t-test, p-value: 0.005725). **(E)** Count of subunits (upper part) and protein complexes (lower part) included in CORUM. Subunits were grouped into SLC, TM-protein and no-TM protein and were counted across all reported protein complexes, taking into consideration multiple occurrences of subunits. **(F)** Distribution of SLC3A2 across the SLC-interactome. The left panel shows the log2 transformed SPC for each SLC AP-MS experiment separated by scored interactions (blue) and background/ not interacting (grey). On the right side it is indicated how often SLC3A2 was identified, scored or found as background. **(G)** Upper part shows for all interactions of SLC3A2 scored within the SLC-interactome (log2FC against GFP, threshold of log2FC > 1). Lower part shows scores of predicted SLC3A2-SLC complexes (high confidence, pDockQ threshold of > 0.5 indicated by dashed line). Complexes for which the experimental structure was solved are marked with an asterisk (*). Predicted complex structures of interactions were compared against a negative set of SLC-chaperone complexes (unpaired student t-test, p-value=0.0009177). On the right side the model of SLC7A7-SLC3A2 is shown. **(H)** SLC-CHP1 interactions in the SLC-interactome (teal). Na+/H+ antiporters of the SLC9A-family are highlighted in purple, other SLCs are grouped by family and coloured in yellow. **(I)** Log2FC against GFP and pDockQ-score of structural models for each SLC-CHP1 complex. Experimentally solved structures are marked in bold and with an asterisk (*). For CHP1 interactions with SLC9A-family members (purple) high confidence models (dashed black line pDockQ > 0.5) were obtained, whereas for the other CHP1-SLC interaction (yellow) only medium (dashed grey line pDockQ > 0.25) to low confidence structures were found. A comparison against randomly sampled SLC-CHP1 interaction showed, a significant difference between models of the interactions covered in the SLC-interactome and the control set (unpaired student t-test, p-value=0.03642). The model of SLC9A2-CHP1 is shown on the right side. For all boxplots in the figure panels: Box represents the interquartile range and its whiskers 1.5 X IQR. Black line represents the median and the black dots represents single measurements.

**Figure EV2.**
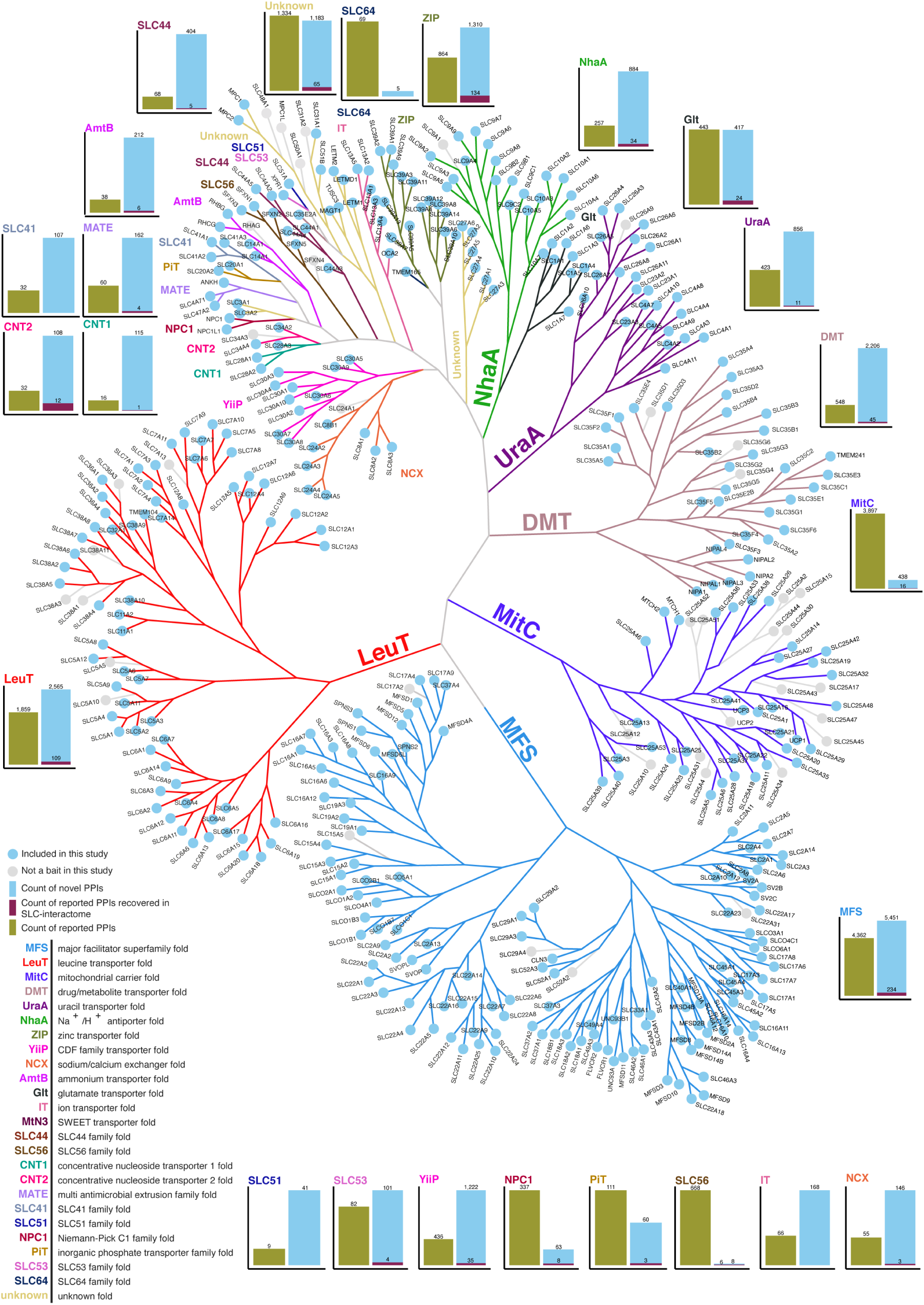
**Novel versus known protein interactions for each clade of the structural and evolutionary based SLCome**. For the construction of the phyologenetic tree the distance matrix of a previous classification of the SLC superfamily based on structural models was used (Ferrada & Superti-Furga, 2022). Each of the 25 distinct structural clades are represented in a different colour. For each clade, the count of novel PPIs (light blue), literature mined PPIs (olive), and the shared PPIs (dark red) are reported. The 405 SLCs included in the study are coloured light blue and all SLCs which were not included in the SLC-interactome are coloured grey.

**Figure EV3.**
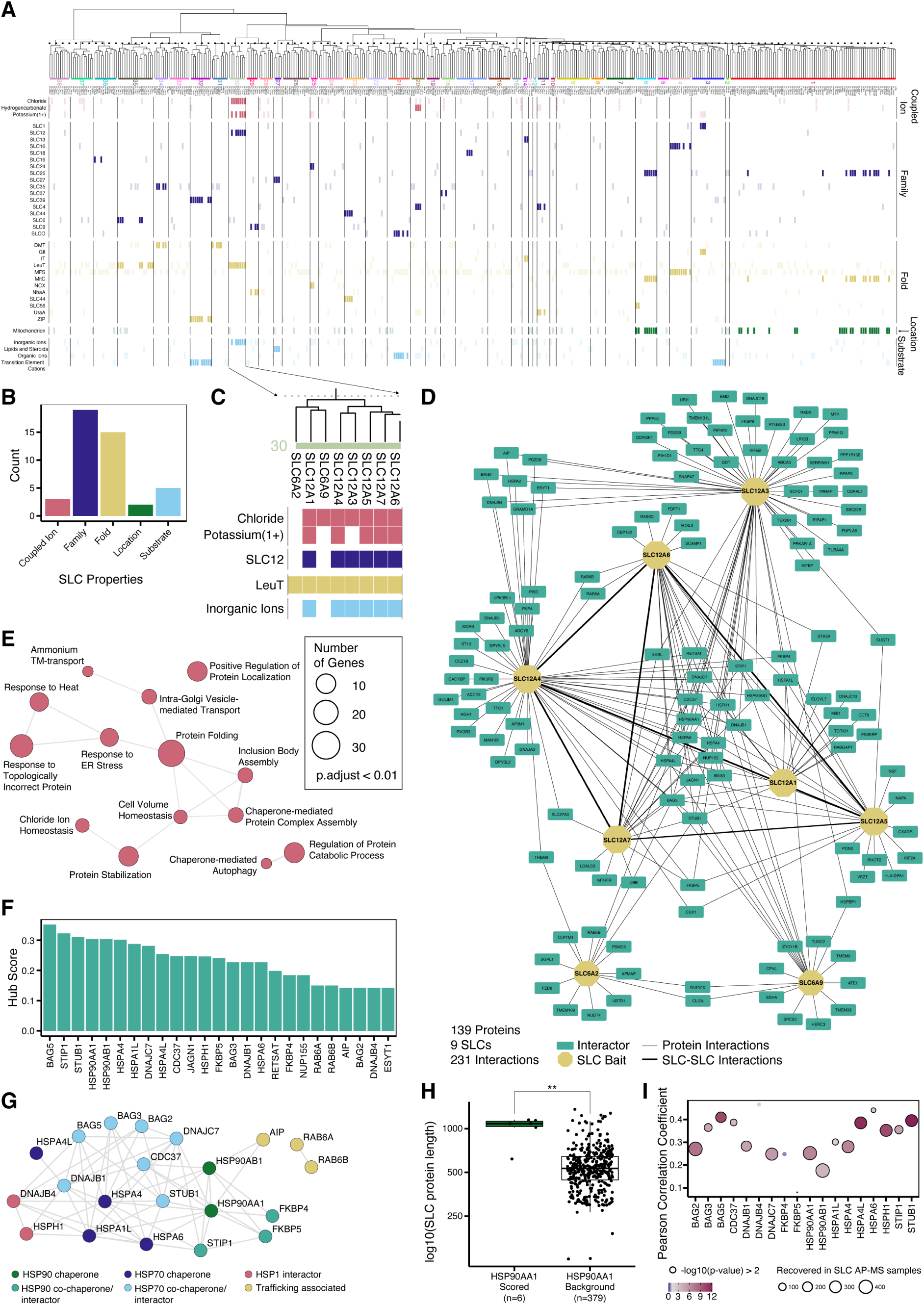
Functional property enrichment analysis showed SLC-interactome similarity for SLC12 and SLC6 family members. **(A)** Significantly enriched SLC functional properties identified in the interactome SLC clustering analysis (Fisher’s test p < 0.2). Colours indicate different SLC functional properties. **(B)** Sum of enriched SLC functional properties separated by SLC properties. **(C)** Significantly enriched SLC functional properties for PPI-interactome profile cluster 30 (Fisher’s test p < 0.2). **(D)** Cluster specific PPI-network obtained by the SLC baits (yellow octagons) grouped to cluster 30. Interactors are shown in green, and interactions between SLC baits are highlighted with a thick black line. **(E)** GO biological processing terms significantly enriched (p-adjusted < 0.01) obtained by GSEA for interactors present in the specific PPI-network. The obtained terms were converted to a similarity network with GO semantic similarity and further filtered to showcase the least overlapping significant terms (similarity cutoff of 0.7). **(F)** The 25 most connected interactors (hub score) in the cluster 30-specific PPI-network. **(G)** PPI-network retrieved from literature (STRING confidence score > 0.4, physical interactions only) for the 25 most connected interactors within cluster 30. Interactors were grouped as HSP90 chaperones (green), HSP90 co-chaperones/interactors (teal), HSP70 chaperones (blue), HSP70 co-chaperones/interactors (cyan), HSP1 interactors (red) or trafficking associated proteins (yellow). The four interactors not connected were removed from the PPI-network. **(H)** Distribution of protein length of SLCs interacting with HSP90AA1 (n=6) and SLCs for which HSP90AA1 was found in the background (n=379). Comparison showed a significant difference in the protein length (student t-test p-value=1.285e-08). **(I)** Correlation of the chaperone/ chaperone interacting protein abundances with the summed SLC tail length (N-terminal and C-terminal). Significant correlations with a p-value < 0.01 are indicated with black ring. For all boxplots in the figure panels: Box represents the interquartile range and its whiskers 1.5 X IQR. Black line represents the median and the black dots represents single measurements.

**Figure EV4.**
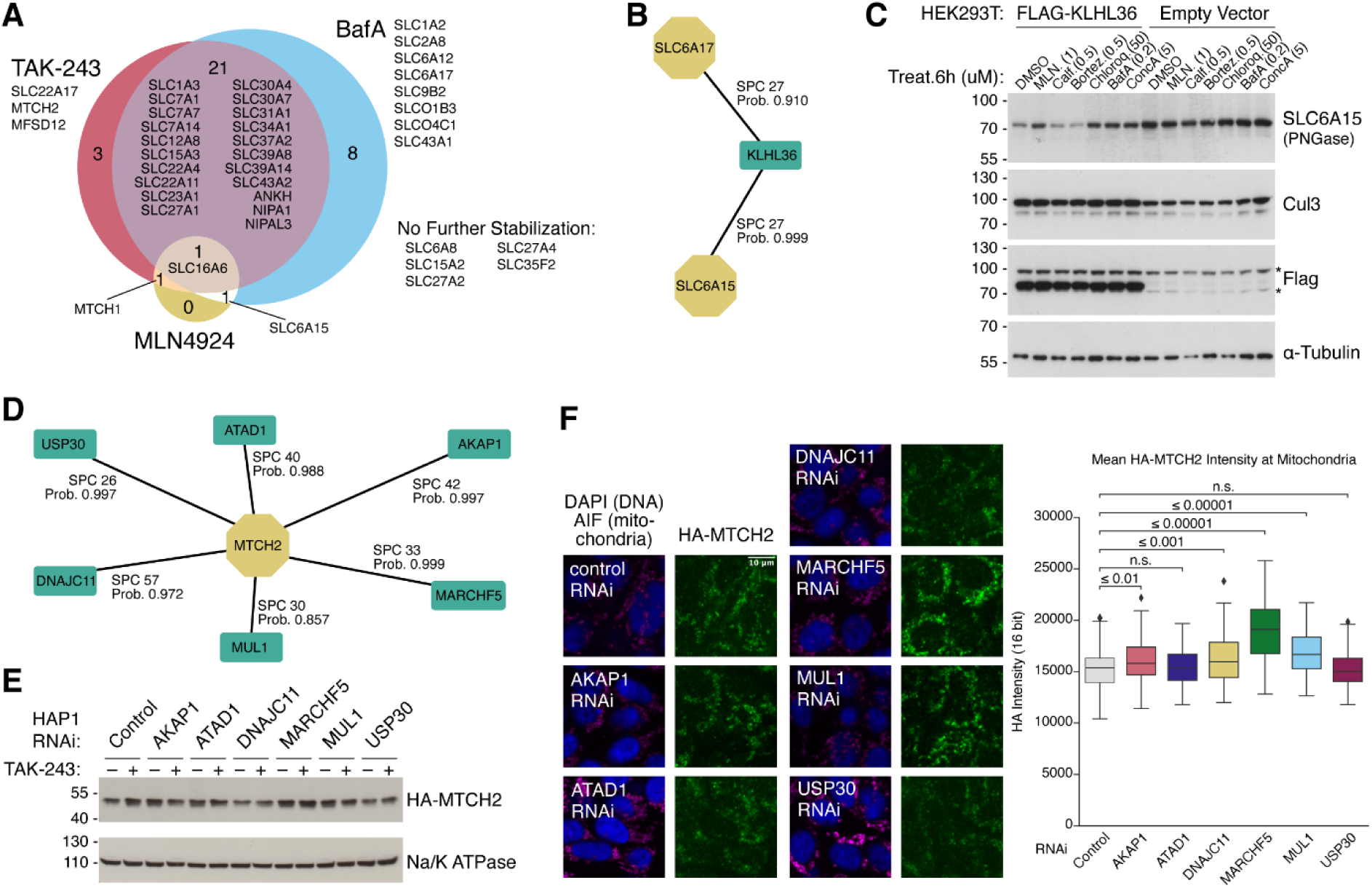
SLC levels are regulated by protein stability. **(A)** Venn diagram of drug treatment effects on SLC protein stability. SLCs were considered to be stabilized by the respective drug if the GFP:RFP ratio was increased more than 10% compared to mock treated cells (p-value < 0.05, independent t test). **(B)** SLC protein interactions of KLHL36. The protein was not quantified in the background of any other SLC purification. **(C)** Western blot result of endogenous SLC6A15 in HEK 293T cells showing SLC6A15 degradation after overexpression of FLAG-tagged KLHL36. **(D)** MTCH2 interactome showing 7 distinct interactors that were assessed in detail. **(E)** HAP1 cells expressing endogenously HA-tagged MTCH2 were transfected with RNAi against MTCH2 interactors. Depletion of the E3 ligase proteins MARCHF5 and MUL1 stabilizes MTCH2 levels as strongly as ubiquitination inhibition by TAK-243. **(F)** Immunofluorescence example images of HA-MTCH2 and quantification at mitochondria as identified by staining for AIF (apoptosis-inducing factor). Over 10,000 cells were imaged per condition; unpaired t-test was used to compare treatments (n = 112 images per sample).

**Figure EV5.**
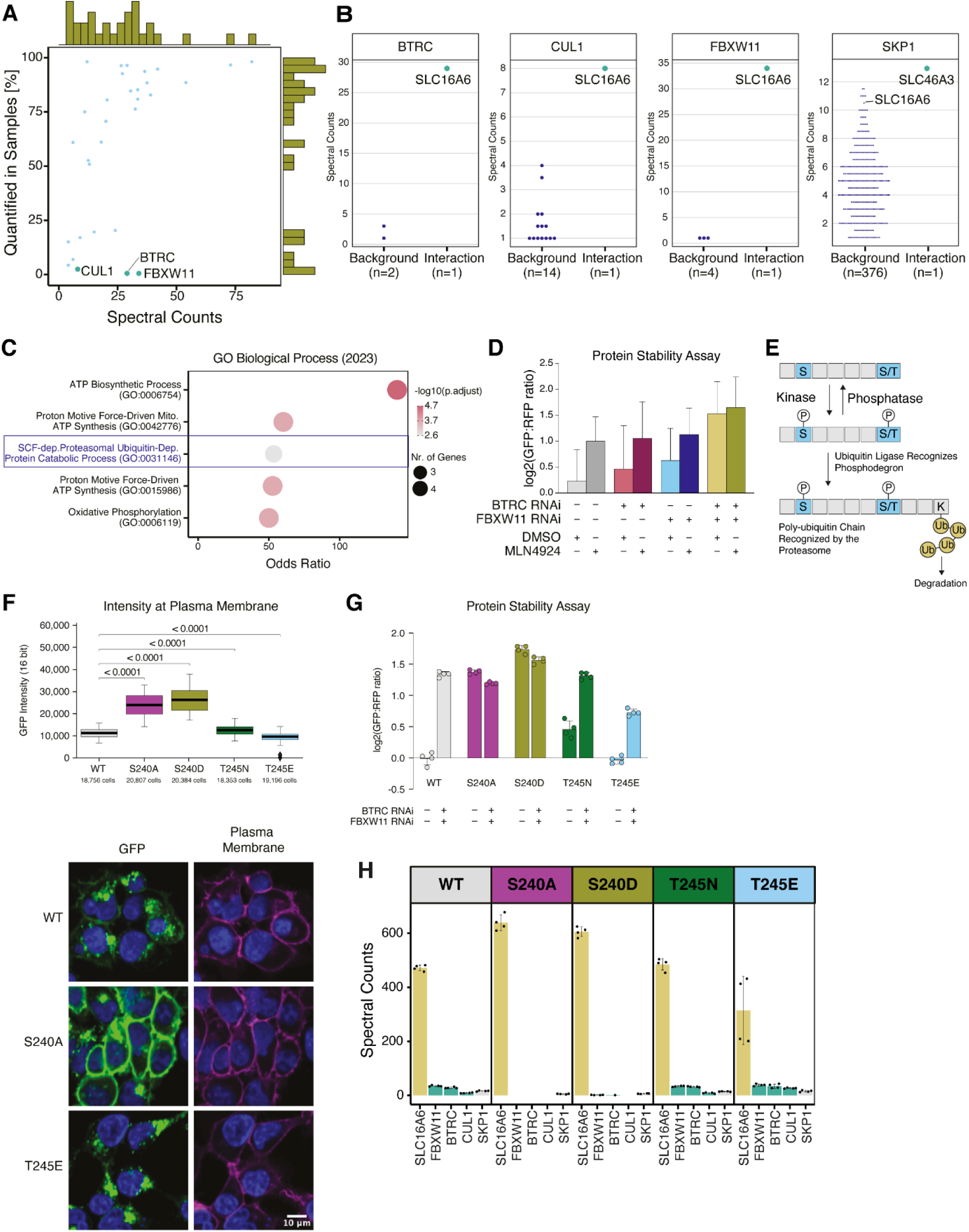
SLC16A6 binds to an SCF E3 ubiquitin-protein ligase complex which mediates abundance. **(A)** Scored interactors (blue points, n=31) of SLC16A6 plotted against identification across the SLC-interactome in percentage. Marginal histograms of data distribution are indicated on the x- and y-axis. SCF complex subunits are labelled. **(B)** Abundance distribution and interactor scoring results of SCF complex members BTRC, CUL1, FBXW11 and SKP1. SKP1 was found in the background of the SLC16A6 purification. **(C)** Five most significant GO biological processing terms enriched in the interactome of SLC16A6. **(D)** Protein stability assay of SLC16A6^WT^ after depletion of BTRC and FBXW11 followed by inhibition of the neddylation pathway by MLN4924. SLC16A6 is fully stabilized by co-depletion of BTRC and FBXW11 without notable stability increase by neddylation inhibition (median + upper quartile, n>14,000 cells per condition). **(E)** Graphical illustration for phospho-degron dependent degradation. **(F)** Quantification of SLC16A6-GFP wild-type and phospho-mutants fluorescence at the plasma membrane (n=160 images per condition; unpaired student t-test was used to compare phospho-mutants against wildtype). **(G)** Protein stability assay of SLC16A6^WT^ and SLC16A6 phospho-mutants in combination with RNAi treatment of adaptor proteins BTRC and FBXW11 (n=4). **(H)** Spectral counts of SCF-complex subunits quantified in AP-MS experiments of SLC16A6 phospho-mutants versus SLC16A6^WT^. Data are represented as mean spectral counts ± SD. Points represent SPC for four MS-injections (n=2 biologically independent replicates with n=2 technical injections).

**Figure EV6.**
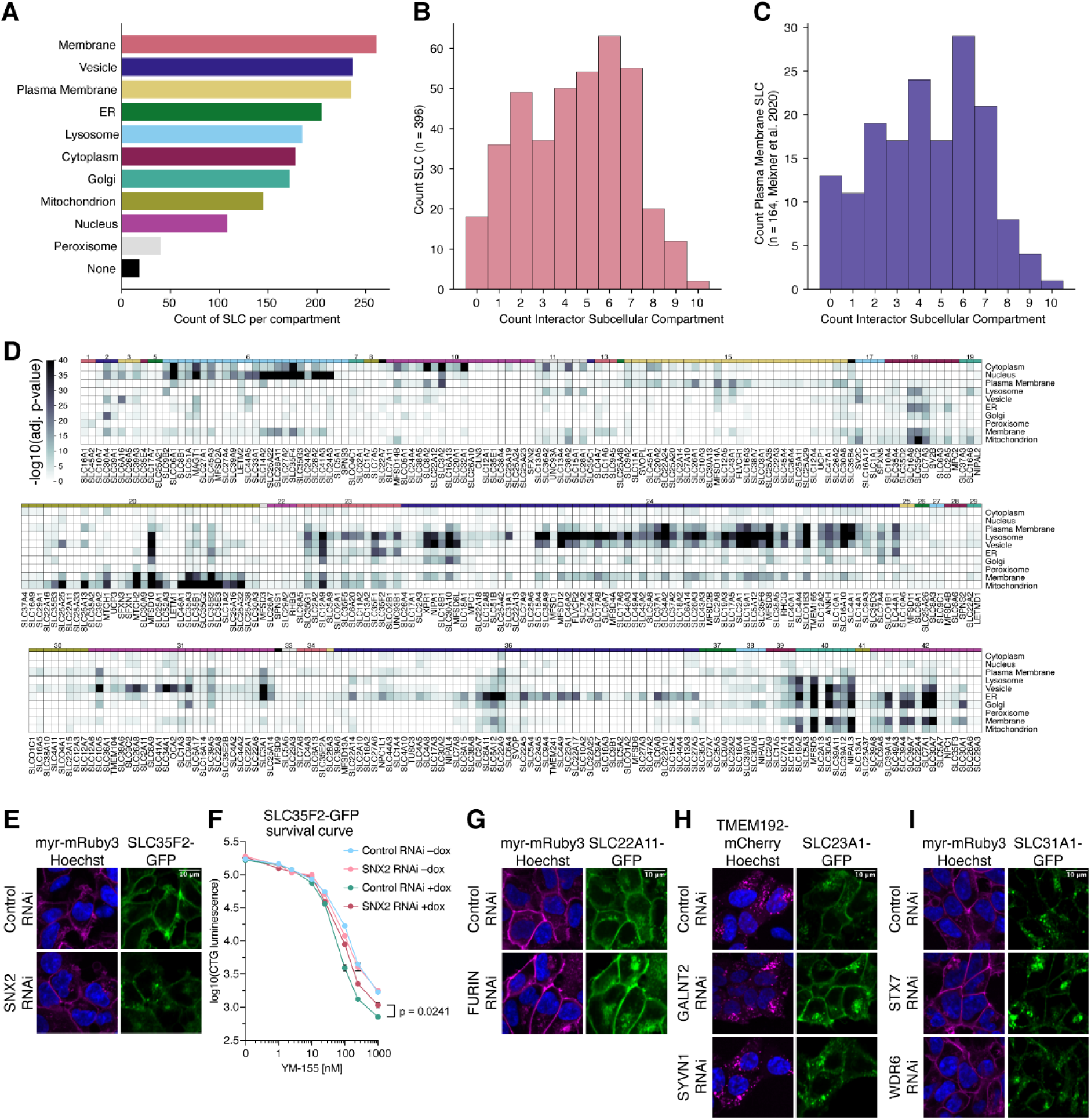
Proteins affecting trafficking and subcellular localization of SLCs. **(A)** GSEA of cellular compartments was performed for each SLC interactome and the resulting GO terms were hierarchically assigned to 10 main compartments (p-value cutoff 0.05; methods for details). **(B)** A majority of SLC interactomes contain interaction partners from several subcellular compartments (mean: 4.5 compartments per SLC). **(C)** This notion also holds true for a subset of SLCs described previously as uniquely plasma membrane-associated (mean: 4.3 compartments). **(D)** The results from the interactome– compartment enrichment analysis were used to cluster SLCs. Strong clusters can be identified for some mitochondria baits (e.g. MTCH2), lysosomal baits (e.g. MFSD12) or Golgi baits (e.g. SLC30A5). **(E)** Representative images of plasma membrane- and Golgi-associated SLC35F2 after RNAi (see Figure 6D for a quantification of the effect; size bar, 10 µm for all images). **(F)** SNX2 depletion attenuates SLC35F2-mediated import of YM-155. After 48 h RNAi, SLC35F2-GFP expression was induced using doxycycline and cells were subjected to YM-155 at the indicated concentrations for 24 h. Cell viability was measured using CellTiter-Glo luminescence (n=3, mean ± SEM). Induced conditions at 1 µM YM-155 were compared using unpaired t test. **(G)** Representative images of plasma membrane- and Golgi-associated SLC22A11-GFP. Depletion of endopeptidase FURIN increased the plasma membrane-associated GFP intensity but not the Golgi-associated GFP intensity. **(H)** Representative images of plasma membrane- and lysosome-associated SLC23A1-GFP. Whereas depletion of GalNAc transferase GALNT2 increased GFP signals overall, depletion of ERAD E3 ubiquitin ligase SYVN1 led to a pronounced increase of GFP overlapping with the lysosomal RFP reference (see Figure 6D for quantifications). **(I)** Representative images of plasma membrane- and vesicle-associated SLC31A1-GFP. Depletion of syntaxin-7 increased overall GFP signal while depletion of WDR6 increased GFP intensity at the plasma membrane (see Figure 6F for quantifications).

